# Mid-cell migration of the chromosomal terminus is coupled to origin segregation in *Escherichia coli*

**DOI:** 10.1101/2023.03.24.534095

**Authors:** Ismath Sadhir, Seán M. Murray

## Abstract

Bacterial chromosomes are dynamically and spatially organised within cells. In slow-growing *Escherichia coli*, the chromosomal terminus is initially located at the new pole and must therefore migrate to midcell during replication to reproduce the same pattern in the daughter cells. Here, we use high-throughput time-lapse microscopy to quantify this transition, its timing and its relationship to chromosome segregation. We find that terminus centralisation is a rapid discrete event that occurs ~25 min after initial separation of duplicated origins and ~50 min before the onset of bulk nucleoid segregation but with substantial variation between cells. Despite this variation, its movement is tightly coincident with the completion of origin segregation, even in the absence of its linkage to the divisome, suggesting a coupling between these two events. Indeed, we find that terminus centralisation does not occur if origin segregation away from mid-cell is disrupted, which results in daughter cells having an inverted chromosome organisation. Overall, our study quantifies the choreography of origin-terminus positioning and identifies an unexplored connection between these loci, furthering our understanding of chromosome segregation in this bacterium.

## Introduction

The faithful and timely segregation of the replicated chromosomes is an essential step in every bacterial cell cycle. While in some bacteria this can be directly attributed to the well-studied ParABS partitioning system, in other species this system is either not strictly essential or is absent altogether. In particular, the mechanism of chromosome segregation in the model system *Escherichia coli* has yet to be identified.

Whatever the mechanism, segregation of the chromosome very likely goes hand in hand with its organisation. Indeed, rod-shaped bacteria have their chromosome arranged linearly within the cell such that the position of each chromosomal locus can be predicted from its genomic position (David et al., 2014; Vallet-Gely and Boccard, 2013; Viollier et al., 2004; Wiggins et al., 2010). However, the orientation within the cell can differ between species and conditions. Typified by C. crescentus, the longitudinal arrangement has the replication origin (*ori*) and terminus (*ter*) located at opposite ends of the cell with the two chromosomal arms organised linearly along the long cell axis (Böhm et al., 2017; Teleman et al., 1998; Vallet-Gely and Boccard, 2013; Viollier et al., 2004). On the other hand, in slow-growing *E. coli* cells, the unreplicated chromosome is believed to adopt a transverse organisation in which the origin (*ori*) is positioned at mid-cell with the left and right chromosomal arms on either side and a stretched terminus region between them (Nielsen et al., 2006b; Wang et al., 2006). Upon replication, the duplicated *ori* segregate outward to the quarter positions with the other replicated loci following progressively, resulting in each replicated chromosome having the same left-*ori*-right organisation and therefore inheritance of the pattern by the daughter cells. However, the symmetry of the above pattern is initially broken by the terminus, which in newborn cells is located close to the new pole i.e. not close to *ori* as might be expected for a circular chromosome. Inheritance of the birth pattern is then achieved by the migration of *ter* to midcell (*ter* centralisation) during the cell cycle (Lau et al., 2003; Nielsen et al., 2006a).

The *ter* sits within the 800 kb Ter macrodomain defined in part by the presence of 23 *matS* sites (Dupaigne et al., 2012; Mercier et al., 2008). The protein MatP binds to these sites and displaces MukBEF, the *E. coli* Structural Maintenance of Chromosomes complex, resulting in a decrease of long-range chromosomal contacts within the region, consistent with its less condensed organisation (Lioy et al., 2018). MatP has been shown to bridge DNA at *matS* sites (Dupaigne et al., 2012). However this is easily out-competed by non-specific DNA binding and so may not be relevant *in vivo* (Crozat et al., 2020; Lioy et al., 2018).

As described above, *ter* is initially located near the new pole but moves to mid-cell during chromosome replication. Its maintenance there is believed to be due to a protein linkage connecting MatP to the divisome protein FtsZ (Espéli et al., 2012; Männik and Bailey, 2015) which facilitates the resolution of chromosome dimers by the FtsK/XerCD machinery just before cell division (Crozat et al., 2020; Stouf et al., 2013). It may also allow *ter* to act as a positive regulator of divisome positioning (Bailey et al., 2014). Despite these studies, the cause of *ter* centralisation and its role within the segregation process has remained unclear.

Here, we use high-throughput single-cell imaging and analysis to quantitatively establish the choreography and timing of *ori* and *ter* in slow-growing *E. coli* cells. We find that *ter* migration from the new pole to mid-cell occurs ~25 min after initial separation of replicated *ori* and independently of its linkage to the divisome. The movement is rapid, occurring within ~5 min, and is coincident with the arrival of *ori* to the quarter-cell positions suggesting a coupling between these two events. Consistent with this, *ter* is unable to stably localise at mid-cell in cells with impaired origin segregation. Overall then, our results show that the mid-cell positioning of the terminus region is not due its replication by mid-cell proximal replisomes or its linkage to the divisome but is rather due to an previously uncharacterized coupling to origin segregation.

## Results

### The cell cycle dynamics of *ori* and *ter*

The accurate analysis of *ori* and *ter* dynamics requires the temporal imaging of a large number of cell cycles. We achieved this using a high-throughput single-cell approach based on a ‘mother-machine’ microfluidic device (Köhler et al., 2022; Wang et al., 2010). Together with a custom analysis pipeline (Köhler et al., 2022), this allowed us to segment and track tens of thousands of cell cycles in steady-state conditions (Fig. 1A, Fig. S1A). To visualize the *ori* region, we used the P1 ParABS labeling system (consisting of *parS*_P1_ inserted near *oriC* and an mTurquoise2-ParB_P1_ fusion expressed from a plasmid (Li et al., 2002; Nielsen et al., 2006a)). The terminus was visualised using a MatP-YPet fusion expressed from its endogenous locus, which has been shown to colocalise well with markers of the terminus region (Bailey et al., 2014; Espéli et al., 2012; Männik et al., 2016; Mercier et al., 2008). Cells grew in the device with a mean cell cycle duration of 133 minutes and mean birth length of 1.71 μm (Fig. S1B,E) and under these conditions we were able to image the cells every 5 minutes while maintaining sufficient signal and without significant changes in growth rate (Fig. S1D).

**Figure 1.**
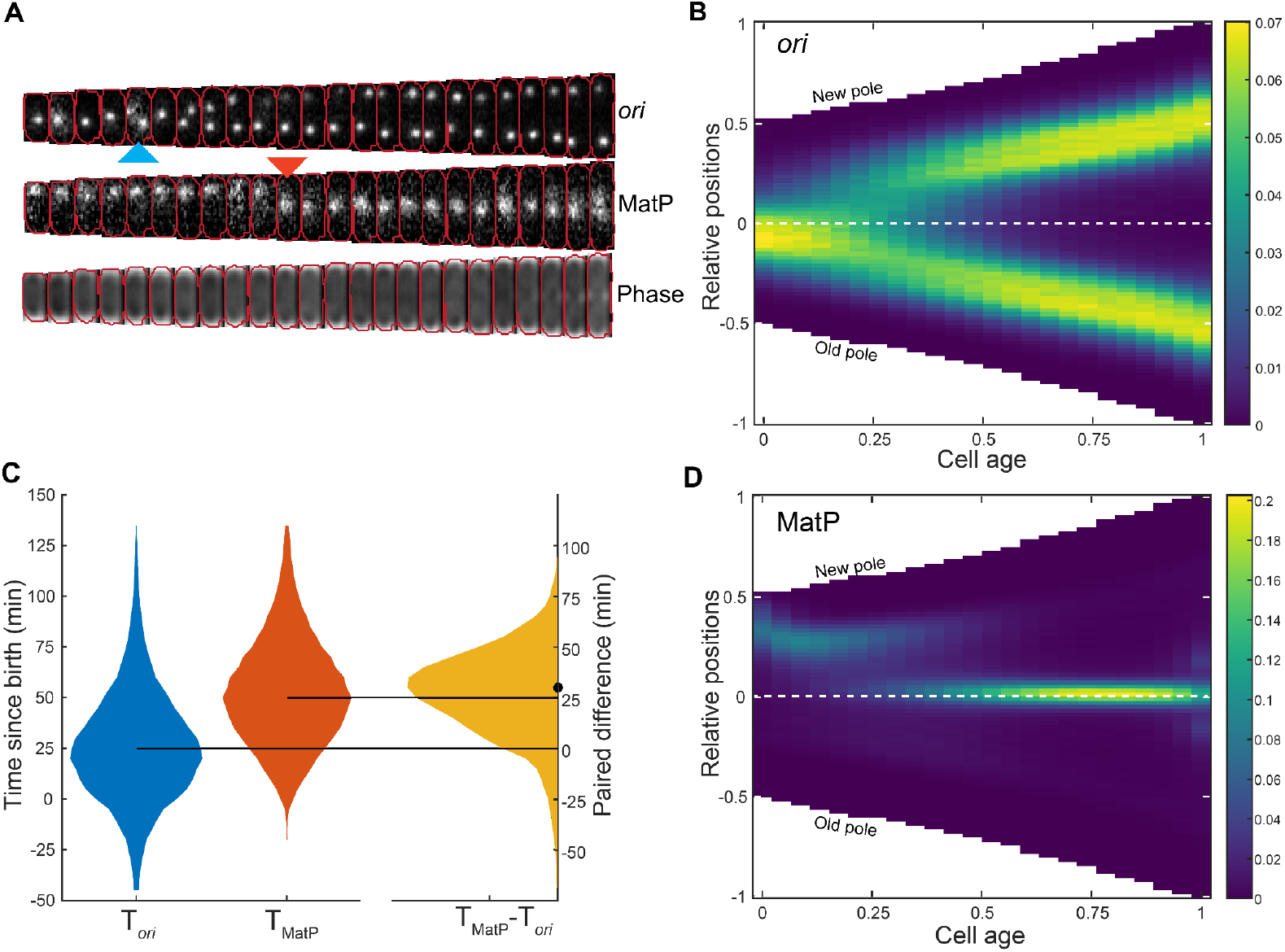
MatP-labelled *ter* re-localizes to mid-cell after *ori* focus duplication. **A.** An example cell cycle with *ori* (ParB_P1_-mTurquoise2) duplication and *ter* (MatP-YPet) relocalization frames marked with blue and red arrows respectively **B.** Population kymograph of *ori* foci positions along the long axis of the cell. Data from different cell cycles are combined by using cell age (0 is birth, 1 is division) and positions relative to an exponentially increasing normalised length. **C.** Distribution of the time of *ori* focus duplication (mean ± sd = 26.6 ± 28.0 min) and MatP relocalization (52.8 ± 26.1 min) along with the time difference between the two events (26.5 ± 23.2 min). The negative values in T*_ori_* and T_MatP_ correspond to events occurring in the previous cell cycle. The horizontal lines indicate the median values of 25 and 50 minutes for T*_ori_* and T_MatP_ respectively. The dot indicates the median (30 minutes) of the paired difference T_MatP_ - T*_ori_*. **D.** Population kymograph of foci positions of MatP positions as in **B.** The values in the color scale for the kymographs represent the frequency of occurrence of foci positions normalized to the number of cell cycles at each cell age. Data is from 38066 cell cycles.

We first confirmed the cell cycle dynamics of *ori*. Consistent with previous studies using agarose pads at similar growth rates (Lau et al., 2003; Li et al., 2002; Nielsen et al., 2006a; Wang et al., 2005), newborn cells typically had a single *ori* focus close to midcell that, upon duplication, segregated outwards to the quarter positions (Fig. 1B). The high-throughput and temporal nature of our data allows us to quantify these observations in detail. Separation of *ori* (defined by two *ori* foci seen for the first time in the cell cycle) occurred on average 27 min after birth, at a cell length of 2.0 μm but with substantial variation between cells (Fig. 1C, Fig. S1L). Indeed, and surprisingly given the slow growth rate, we found that 15% of cells were born with more than one *ori* focus (Fig. S1F), indicating that DNA replication was initiated in the previous cell cycle (note that this is not detectable in the population average kymographs (Fig. 1B) or demographs (Fig. S1G). We have shown elsewhere that this is consistent with the volume dependence of chromosome replication initiation (Donachie, 1968; Levin and Taheri-Araghi, 2019) and arises from a second replication initiation in the mother cell due to the size of the mother cell crossing the volume per origin initiation threshold for a second time in the same cell cycle (Köhler et al., 2023).

We also confirmed the positioning of the *ter* region - it was found close to the new pole at birth before moving to midcell, where it remained tightly localised for the majority of the cell cycle (Fig. 1D). To quantify the timing of this transition, we defined *ter* arrival at midcell as the first frame at which the MatP-YPet focus is within the middle 4.8 pixels (320 nm) of the cell for 3 consecutive frames, the former value being determined from the position distribution (Fig. S1K). Using this measure, stable *ter* localisation to mid-cell occurred on average 53 min into the cell cycle and 26 min after *ori* separation (Fig. 1C). These timings are consistent with those previously inferred from snapshots (Wang et al., 2005). Here, however, we follow complete cell cycles and have captured the entire distribution of timings.

The visible separation of *ter* foci occurred just before cell division consistent with its association to the early divisome protein FtsZ, though the timing of this relative to the cell cycle is dependent on our identification of the cell division event. Correspondingly, at birth *ter* foci were initially found closer to the pole before moving inward to the edge of the nucleoid (Fig. 1A,D and below).

### Origins and nucleoid are asymmetrically positioned

Our data also show that *ori* is not precisely positioned at the mid- and quarter-cell positions but rather exhibits a bias towards the old pole at birth and to either pole before division. The offset is small (about 5% of cell length) but reproducible and persistent during the beginning of the cell cycle. As a consequence, the trajectories of segregating *ori* foci are not symmetric with the new-pole proximal *ori* moving further and faster (see below) to reach its target quarter cell position. Note that the bias at birth is only apparent when cells are ordered according to their polarity. It is not detectable when cells are oriented randomly, as would be the case for a snapshot-based analysis (Fig. S2A). The mid-cell positioning of *ter*, on the other hand, is precise (Fig. 1D).

Since the nucleoid exhibits a new-pole bias during the early part of the cell cycle (Bates and Kleckner, 2005; Fisher et al., 2013; Hadizadeh Yazdi et al., 2012), we sought to determine how *ori* is positioned relative to the nucleoid. We therefore examined the localisation of *ori* and *ter* in a strain expressing the nucleoid marker HU-mCherry (Fig. 2A). Consistent with previous results, we found a clear bias of the nucleoid toward the new pole that gradually decreases during the first half of the cell cycle until the nucleoid is symmetrically positioned within the cell (Fig. 2B). As a result, at birth the *ori* is positioned at the old-pole proximal periphery of the nucleoid, typically at the outer quarter mark of (background-subtracted) HU-mCherry signal. After duplication, one *ori* moves to the opposite side of the nucleoid resulting in a symmetric configuration both with respect to the nucleoid and the cell (Fig. 2C). Interestingly, the position of *ter* is unaffected by the initial bias in the nucleoid position, perhaps because the bias has largely been resolved by the time of *ter* centralisation (Fig. 2D).

**Figure 2.**
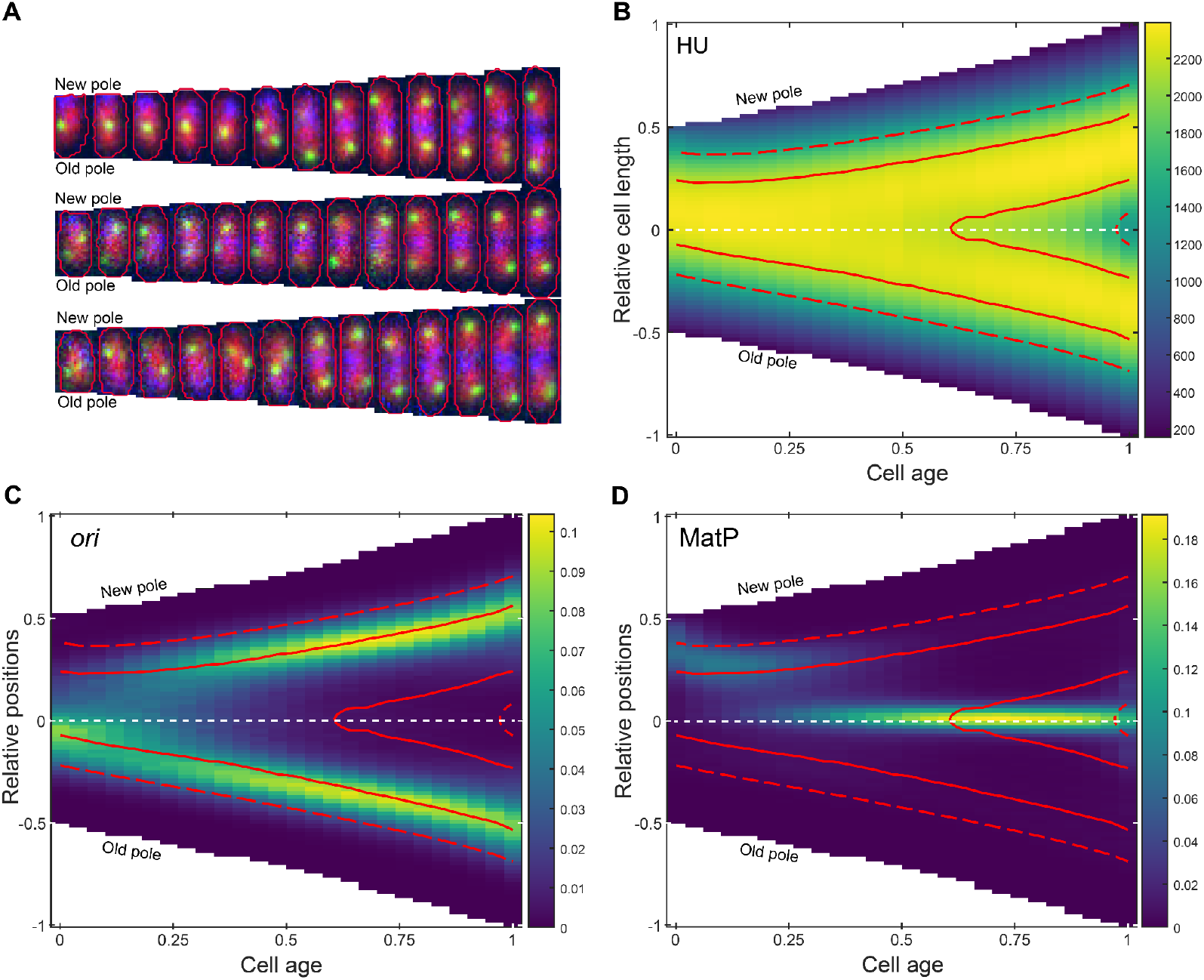
The *ori* is positioned at the periphery of the nucleoid. **A.** Representative cell cycles (10-minute intervals) with *ori* in green, HU in red and MatP in blue **B.** Population kymograph of HU-mCherry signal along the long axis of the cell. The solid and dashed contour lines enclose the upper 50 and 80 percent respectively of the total HU-mCherry signal. **C.** Population kymograph of *ori* foci positions (as in Figure **1B**) with contour lines from **B**. **D.** Population kymograph of MatP foci positions (as in Figure **1D**) with contour lines from **B.** The values in the colormap scale for **B** represents the mean intensity of line profiles across the cell length. The color scale in **C** and **D** is as in Figure 1. Data is from 33593 cell cycles.

Overall these results refine our understanding of *ori* and *ter* positioning in slow-growing cells. In terms of positioning within the cell, our analysis reproduces the established view, largely based on snapshot imaging (Fekete and Chattoraj, 2004; Lau et al., 2003; Li et al., 2002; Nielsen et al., 2006a; Wang et al., 2005), that *ori* exhibits mid-cell positioning at birth, albeit with the addition of a slight old-pole bias, followed by segregation to the quarter positions. However, it is also generally assumed that *ori* is centrally positioned within the nucleoid consistent with an observed left-*ori*-right chromosome organisation (Nielsen et al., 2006b; Wang et al., 2006). Our results do not support this assumption - the *ori* lies toward the nucleoid periphery both before initiation at the beginning of the cell cycle and after segregation. This agrees with earlier studies using FISH and DAPI-labelling (Bates and Kleckner, 2005; Fisher et al., 2013; Niki et al., 2000; Niki and Hiraga, 1998). Thus, the current model of a nucleoid-centered left-*ori*-right configuration during slow growth does not hold in its entirety and may bely the complexity of chromosome organisation in this bacterium (see discussion below).

### *ter* centralisation occurs before nucleoid constriction

It was recently suggested that *ter* centralisation shortly precedes the stable appearance of a constricted/bilobed nucleoid structure (Männik et al., 2016), which limits the onset of cell division (Tiruvadi-Krishnan et al., 2022). To examine this, we analysed the HU-mCherry signal across the long-axis of the cell and identified the time of stable (i.e. not transient) nucleoid constriction (Fig. 3A, S3A,B). In contrast to the results of Männik et al, we found that stable constriction occurs significantly later (45 min) than *ter* centralisation (Fig. 3B), suggesting no direct causal relationship between these two events. Nevertheless, we have now identified the timing of three important cell cycle events - separation of duplicated origins, *ter* centralisation and the onset of nucleoid constriction (Fig. S3C). While there is significant variation in the timing of these events between cells, in almost all cells they occur in the given order (Fig. S3D).

**Figure 3:**
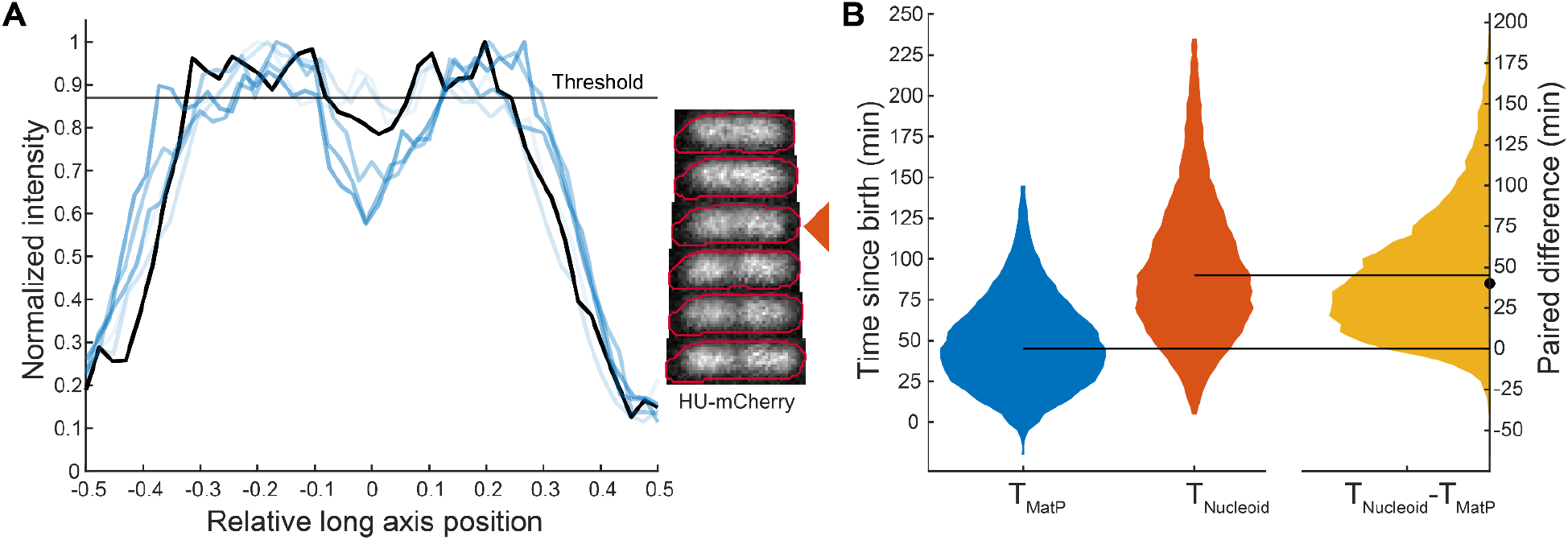
MatP relocalization precedes nucleoid constriction. **A.** Line profile plots of HU-mCherry signal on different frames with corresponding images. The nucleoid is considered constricted if the relative depth of the dip in the signal (if one exists) is greater than a threshold chosen to account for normal variation in the signal (see Fig. S3A,B). The threshold for the frame indicated by the arrow is shown in gray. The time of stable nucleoid constriction T_Nucleoid_ is the earliest frame after which a dip greater than the threshold is observed in the nucleoid signal for the rest of the cell cycle. **B.** Distribution of the time of MatP centralization, T_MatP_ (mean ± sd = 49.5 ± 28.4 min) and stable nucleoid constriction, T_Nucleoid_ (94.2 ± 43.7 min) along with the time difference between the two events (51.5 ± 38.1 min) as in Figure 1. The horizontal lines indicate the median values of 45 and 90 minutes for T_MatP_ and T_Nucleoid_ respectively. The dot indicates the median (40 minutes) of the paired difference T_Nucleoid_ - T_MatP_. Data as in Figure 2. See also Fig. S2 and S3.

### *ter* centralisation coincides with completion of *ori* segregation

The kymographs and demographics of MatP positions (Fig. 1C, Fig. S1G) reveal separated peaks, indicating that the migration of *ter* from the edge of the nucleoid to midcell is relatively rapid compared to its movement at other times in the cell cycle. In fact, this transition could occur within a single frame (5 min) (Fig. 1A). This could be made apparent by synchronizing the cell cycles according to the time of *ter* centralisation (Fig. 4A). Furthermore, the mean step-wise velocity of MatP foci (measured between consecutive frames) sharply peaked at the transition before dropping rapidly to zero afterwards (Fig. 4B). Notably, this is not a consequence of the synchronization, as the greatest movement most frequently occurred at the transition (Fig. S4).

**Figure 4:**
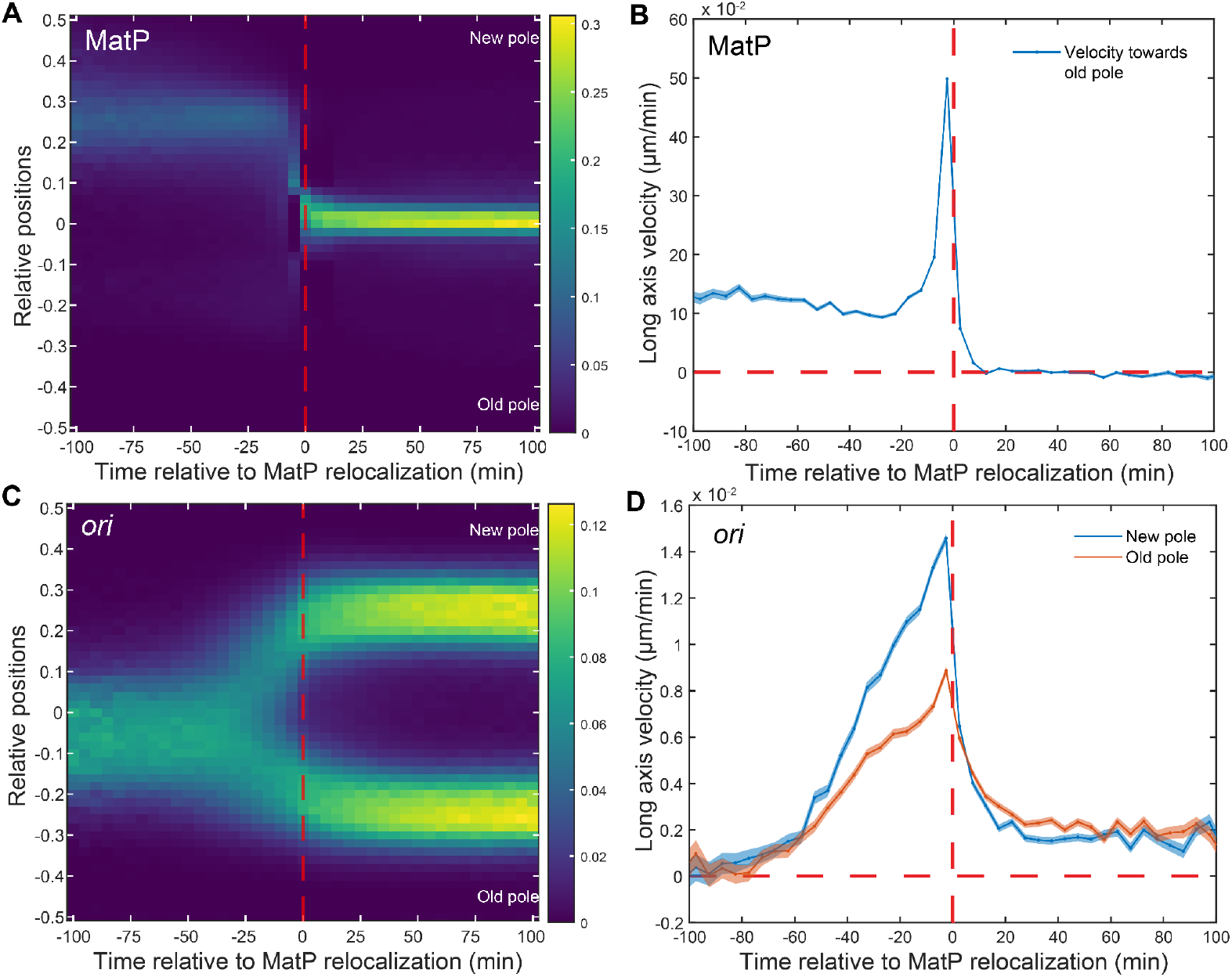
*ter* centralisation coincides with completion of *ori* segregation. **A.** Kymograph of MatP foci positions synchronized to MatP centralisation. **B.** Mean velocity of the MatP focus towards the old pole relative to the time of MatP centralisation. **C.** Kymograph of *ori* foci positions synchronized to MatP centralisation **D.** Mean velocity of *ori* foci towards the nearest pole relative to the time of MatP centralisation. Data as in Figure 1. The color scale in **A** and **C** is as in Figure 1. Shading in **B** and **D** indicate standard error of the mean.

While the migration of *ter* to mid-cell has previously been attributed to the action of the mid-cell localized replication machinery (Espéli et al., 2012), it is unclear if this is consistent with such a rapid transition. Indeed a study of chromosome organization during fast growth found that *ter* centralisation was correlated with cell length rather than progression of the replication fork and that, irrespective of when the transition occurred the remaining unreplicated DNA migrates inwards (Youngren et al., 2014). This is consistent with the large variation we observe in the timing of the transition (Fig. 1C), which can occur even before visible origin separation or as late as 75 min afterwards.

On the other hand, the kymographs in Figure 1 indicate that *ter* centralisation occurs at approximately the same time as *ori* segregation. In fact, after synchronizing the *ori* foci positions relative to *ter* centralisation, it became clear that centralisation is coincident with the completion of *ori* segregation, i.e, with the arrival of the replicated *ori* at the quarter positions of the cell (Fig. 4C). The average velocity of both the new pole and old-proximal *ori* increased steadily up to the *ter* transition before dropping rapidly, with the peak occurring at the same time as that of Ter (Fig. 4D). We additionally note that the new pole-proximal *ori* exhibits a higher mean velocity than its sister, consistent with our previous observation of asymmetric ori segregation (Figure 2). Overall these results indicate a coupling between *ter* centralization and the completion of origin segregation and it is tempting to speculate a causative relationship between the two events, namely that the final stage of *ori* segregation somehow triggers *ter* to rapidly move to midcell.

### *ter-ori* coupling does not depend on the *ter* linkage

If *ter* centralisation and *ori* segregation are genuinely coupled, then disrupting one or the other process may provide insight into their codependency. In this direction, we first targeted *ter* centralisation. As discussed above, while the cause of *ter* migration to midcell is unclear, on arrival it is partially anchored to the divisome by a protein linkage involving FtsZ, ZapA, ZapB and MatP (Espéli et al., 2012). Disrupting this linkage has previously been shown to reduce the duration of *ter* centralisation and alter the timing of sister *ter* segregation (Espéli et al., 2012; Männik et al., 2016; Nolivos et al., 2016). However, when we imaged a *zapB* deletion strain, we found that the effect on MatP foci positioning was relatively minor, with only a slight broadening of its position distribution and slightly earlier segregation (compare Fig. S5 and Fig. 1). The transition to mid-cell still occurs rapidly (Fig. 5A,B). Interestingly, the asymmetry in *ori* positioning is almost absent with only a small difference in the velocity of sister *ori* detectable (Fig. 5C,D). Nevertheless, *ter* centralisation is still coincident with completion of origin segregation (Fig. 5B,D).

**Figure 5:**
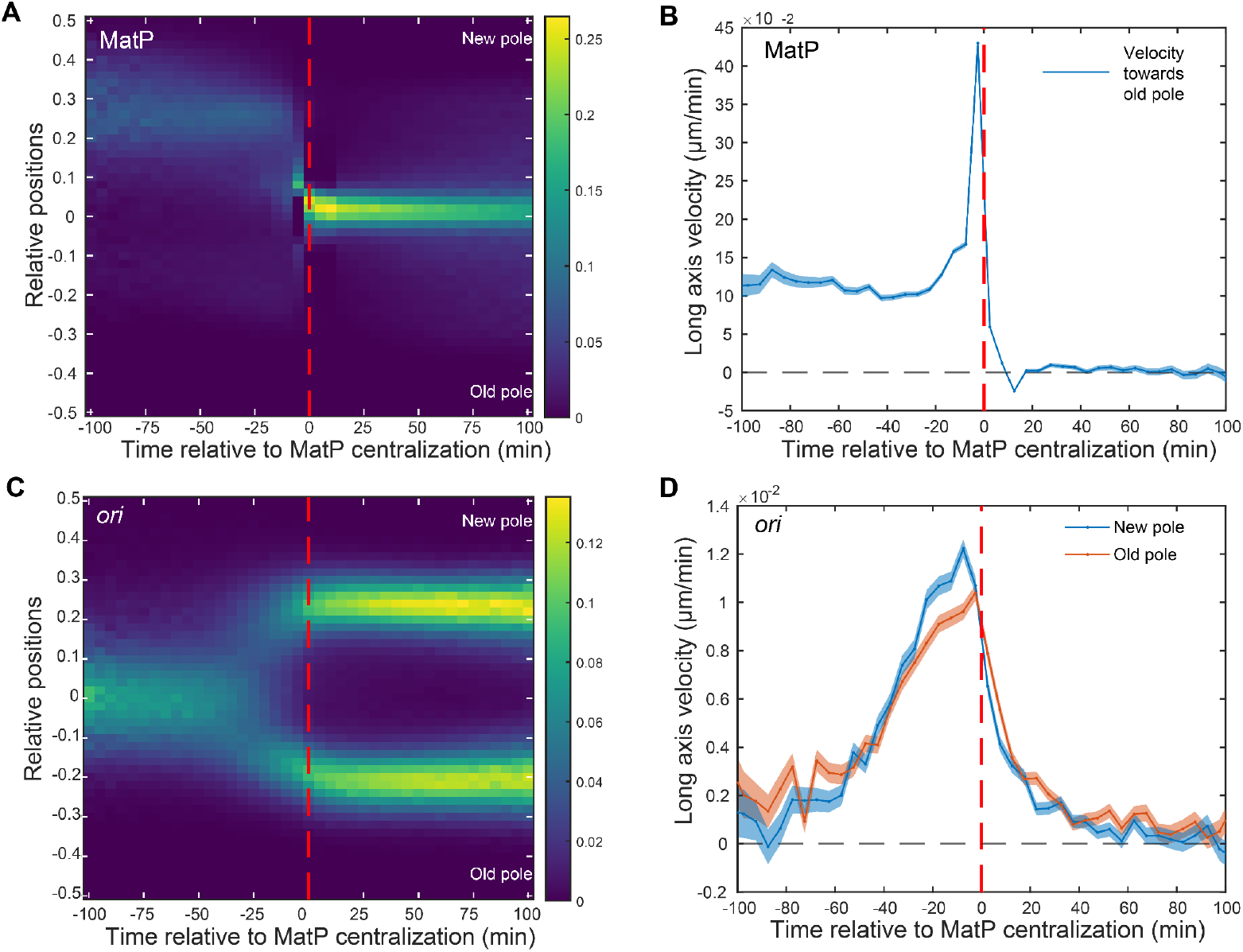
*ter*-linkage is not involved in MatP relocalization. **A.** Kymograph of MatP foci positions in a *zapB* strain synchronized to MatP centralisation. **B.** Mean step-wise velocity of MatP focus tracks towards the old pole relative to MatP centralisation in a *zapB* strain. **C.** Kymograph of *ori* foci positions synchronized to MatP centralisation in a *zapB* strain **D.** Mean velocity of *ori* foci tracks towards the nearest pole relative to the time of MatP centralisation in a *zapB* strain. Data is from 25282 cycles. Shading in **B** and **D** indicate standard error of the mean. The color scale in **A** and **C** is as in Figure 1.

A stronger phenotype was obtained in *matPΔC20* cells, in which MatP lacks 20 amino acids from its C-terminal, which is believed to prevent its multimerisation and interaction with ZapB but not *matS* binding (Crozat et al., 2020; Dupaigne et al., 2012; Espéli et al., 2012; Nolivos et al., 2016). While we again found that the timing of *ter* centralisation is very similar to wildtype, *ter* is more dynamic and often overshoots the mid-cell before returning (Fig. S6A). This was apparent at the population level as a smear in the kymograph (Fig. S6C). The segregation of sister *ter* occurs noticeably earlier and as a result there is a correspondingly shorter period of centralisation (Fig. S6C,E). Despite these abnormalities, *ori* positioning seems largely unaffected (Fig. 6B,F) and importantly the completion of *ori* segregation remained coincident with *ter* centralisation (Fig. 6 E,F,H,I). We also noted that the time between *ori* duplication and *ter* centralization is decreased in both mutants compared to wild-type. As it seems unlikely that these mutations would increase the duration of duplicated *ori* cohesion or accelerate DNA replication, this supports the idea that *ter* migration is not triggered directly by replication fork progression. Overall, these results confirm that the migration of *ter* to mid-cell, and its maintenance there, does not require the linkage of the terminus region to the divisome, while at the same time showing the robustness of the coupling between *ter* centralisation and completion of *ori* segregation.

**Figure 6:**
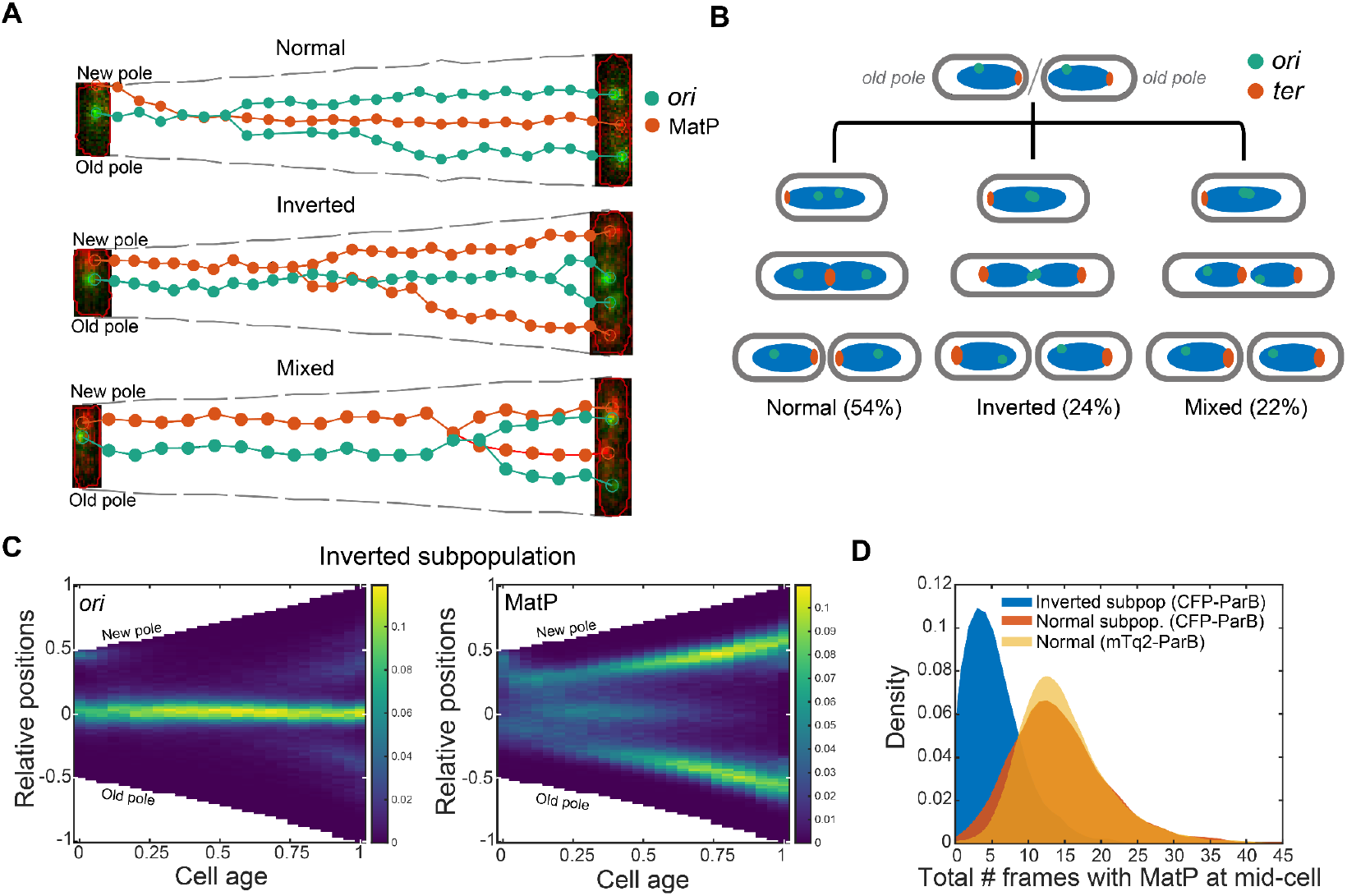
Stable MatP localization at mid-cell requires *ori* segregation. **A.** Example cell cycles resulting in normal, inverted or mixed (one normal, one inverted) orientation of daughter cells. Foci position of *ori* (green) and MatP (red) are shown with a composite fluorescent image of the first and last frame of the cell cycle. **B.** A schematic representation of *ori-ter* organization based on **A** and the previous nucleoid imaging**. C.** Foci position kymographs of the inverted subpopulation (n=3774 cycles) **D.** Distribution of the total number of frames in which MatP foci is found at mid-cell in each cell cycle of the inverted subpopulation (*ori*-CFP-ParB, n=3774 cycles), normal subpopulation (*ori*-CFP-ParB, n=8269 cycles) and for the data from **Figure 1** (*ori-*mTq2-ParB, n=38066 cycles). The color scale in **C** is as in Figure 1.

### *ori* segregation is a requirement for stable MatP relocalization to mid-cell

We had more success disrupting *ori* segregation. The original ParB_P1_ labeling system produces foci numbers consistent with flow cytometry when used at basal expression levels, i.e. uninduced (Nielsen et al., 2006a, 2006b). However, high levels of induction result in a segregation defect with fewer and larger ParB foci detected (Nielsen et al., 2006a). This was an initial challenge as continuous imaging in the mother machine requires sufficiently high expression and continuous induction. However, we found that the defects were attributable to the dimerisation of the CFP fluorophore used since no defects were detectable when using the monomeric mTurquoise2 fusion to visualize *ori*, presented thus far (see methods). Nevertheless, this finding was fortuitous as we can use the induced CFP-ParB_P1_ system as a tool to disrupt *ori* segregation. This has advantages over, for example, depleting TopoIV (Nicolas et al., 2014) as the direct effect of the perturbation should be local to the *ori* region.

We found that more than half of the cells with induced expression of CFP-ParB_P1_ displayed the same spatiotemporal organisation of *ori* and *ter* at division as seen previously for the mTurquoise2-ParB_P1_ labeling system (Fig. 6A, top). However, the remaining ~46% showed a defect in *ori* segregation, with only a single mid-cell localised CFP-ParB_P1_ (*ori*) focus visible for most of the cell cycle (Fig. 6A, middle and bottom). At the same time, *ter* (MatP-YPet) does not maintain a sustained mid-cell localisation as in normal cells. It still moves to mid-cell but only for a short time, likely in order to be replicated, as evidenced by the appearance of two foci shortly afterwards. These MatP foci were often found to rapidly move outward towards opposite poles, resulting in both daughter cells having an inverted *ori-ter* axis i.e. with *ter* at the old rather than new poles (Fig. 6A middle, 6B). Alternatively, one chromosome manages to correct itself before division resulting in only one daughter cell having an inverted orientation (Fig. 6A bottom, 6B). Interestingly, despite these defects, the growth rate and cell cycle duration of the cells were comparable (Fig. S7E,F). This shows that while a new-pole-oriented *ter* is the norm, neither it nor *ori* segregation away from mid-cell, are requirements for successful completion of the cell cycle. Indeed, in our data sets in Figures 1 and 2, we also find cells with an inverted orientation, albeit at a very low frequency of 0.5-1%. We also do not observe a hereditary effect on orientation - cells born with an inverted orientation have a similar probability to produce inverted daughters as cells born with a normal orientation (Fig. S7C).

This mis-positioning of *ori* and *ter* are clearly reflected in the averaged kymographs, which contrast strongly with the normal population (Fig. 6C, Fig. S7A,B). Note that while *ter* shows some period of mid-cell localisation in these kymographs, it is a result of variation between cells: It spends on average one third of the time at mid cell compared to the normal subpopulation or to cells with *ori* labeled using mTurquiouse2-ParB_P1_ (Fig. 6D). Colocalisation with *ori* is also not increased compared to normal cells (Fig. S7D). Overall, these results suggest that segregation of sister *ori* away from mid cell is required before *ter* can be stably localised to that position.

What might underlie this apparent repulsion between *ori* and *ter*? The Structural Maintenance of Chromosomes complex MukBEF is an important chromosomal organiser in *E. coli* and is required for correct *ori* positioning (Badrinarayanan et al., 2012; Danilova et al., 2007; Hofmann et al., 2019). It forms DNA-associated clusters that colocalise with *ori* but is displaced from the terminus region by its interaction with MatP (Lioy et al., 2018; Nolivos et al., 2016), making it a plausible mediator of the coupling we observe. However, we found that *ter* positioning is largely unchanged in the absence of MukBEF (Fig S8A) consistent with a recent study (Mäkelä et al., 2021). Importantly, it still displays a similar rapid transition from the new pole to midcell. Unfortunately, our *ori* labeling system (mTurquoise2-ParB_P1_) appeared to induce defects in the absence of MukBEF as evidenced by a large increase in anucleate cells (occurring in approximately 25% of divisions) and a somewhat disrupted *ter* positioning when we additionally labelled *ori* (Fig. S8B,C). It may be that TopoIV recruitment by MukBEF is required to counteract supercoiling induced by ParB (Lemonnier et al., 2000). We therefore cannot determine if there is a difference in the relative timing of *ter* centralisation and *ori* segregation. However the snapshot demographs of Mäkelä et al. based on FROS labeling of loci, though more limited in their resolution, appear to show that *ori* segregation from mid-cell to the poles is still roughly coincident with *ter* centralisation in the absence of MukBEF. While further work is required, these data suggest that MukBEF is likely not responsible for the coupling we observe. Note that we have no reason to suspect artifacts from the mTurquoise2-ParB_P1_ labeling system in the other strains studied. The foci distributions we obtain are consistent with previous studies and no phenotypic changes were observed when a lower induction level was used.

## Discussion

### *ter* centralisation is coupled to, and requires, *ori* segregation

Previous work based on fluorescence *in-situ* hybridisation (FISH) defined several organisational transitions of the *E. coli* cycle during slow growth (Bates and Kleckner, 2005). The T1 transition is defined by the initial separation of sister *ori* at mid-cell. One sister moves to the *ter*-distal end of the nucleoid while the other remains close mid-cell. The T2 transition occurs ~20 min later and marks the rapid movement of the remaining mid-cell-proximal *ori* to the *ter*-proximal edge of the nucleoid. This is also coincident with the movement of *ter* to mid-cell. The result is that the sister *ori* are positioned symmetrically at opposite ends of the nucleoid which correspond to the quarter positions of the cell. Our results are in broad agreement with this study. However, while we do also observe an asymmetry in *ori* positioning and movement, our data show both sister *ori* reaching their target positions at the edge of the nucleoid simultaneously. Our tracking of tens of thousands of cell cycles at a higher temporal resolution also allow us to establish that *ter* centralisation is much more rapid than *ori* movement, occurring in about 5 min (1 frame) and more specifically is coincident with completion of the more gradual process of *ori* segregation.

What might underlie such a rapid transition? We find it unlikely that it is due to the action of the typically mid-cell localized replisomes given the large variation in the time between initial separation of *ori* foci and *ter* centralisation (Fig. 1C). The same conclusion was made in a study of chromosome organisation during fast growth (Youngren et al., 2014), which found that the transition can take place at very different stages in the replication process but that whenever it occurs any unreplicated DNA is brought with the terminus to mid-cell. This was attributed to the entropic elasticity of a stretched polymer. Indeed, polymer simulations have shown that a partially replicated chromosome in a rod-shaped cell will organise itself to place the unreplicated terminus region at mid-cell (Jun and Mulder, 2006) and this can lead to rapid movement of the terminus region from the pole to mid-cell (on the timescale of the stochastic fluctuations of mean terminus position) (Männik et al., 2016). We speculate that this entropy-driven process could be triggered or promoted by the outward segregation of the replicated *ori*, which we have previously proposed may be due to the action of self-organising MukBEF (Hofmann et al., 2019; Murray and Sourjik, 2017).

### The role of the *ter*-linkage

The *ter* region is believed to be anchored to the septal ring through a MatP-ZapB-ZapA-FtsZ protein linkage. However, disruption of this linkage was found to reduce the duration of, but not entirely abrogate, the mid-cell localisation of *ter* and leads to earlier separation of sister loci (Espéli et al., 2012; Männik et al., 2016). However, while we indeed observed this for the *matPΔC20* strain (Fig. S6), deletion of *zapB* had a very mild effect on *ter* centralisation, with only a slightly wider position distribution and slightly earlier release of sister *ter* detectable (Fig. S5). This suggests that either MatP, through its C-terminal domain, may interact with other components of the divisome or that some part of its anchoring effect requires its C-terminal domain independently of its linkage to the divisome. The latter could be connected to MatP multimerization and bridging of sister *ter* (Dupaigne et al., 2012; Mercier et al., 2008).

### *ori* segregation is not required for bulk chromosome segregation

In *E. coli* and other studied bacteria, chromosome segregation begins with the origin and proceeds progressively as the chromosome is replicated. Indeed, the importance of initial origin segregation is underlined by the presence, in many species, of a dedicated system (ParABS) responsible for this task. It was therefore surprising to find that the origin segregation defect occurring in roughly half of cells using the CFP-ParB labelling system (and ~0.5% of cells using mTurquoise2-ParB), did not prevent or inhibit successful completion of the cell cycle (Fig. 6).

These cells maintained unsegregated *ori* at mid-cell for the majority of cell cycle but nevertheless segregated their chromosomes into each cell half, producing daughter cells with an inverted chromosome orientation i.e. *ter* at the old and *ori* at the new pole. This result indicates that while origin segregation typically leads the process of chromosome segregation, it is not a requirement for it to do so. Note that since we label one specific locus (14 kb from *oriC*), it is possible that the rest of Ori macrodomain segregates properly with the *parS* proximal regions being maintained together at mid-cell. However, this would not explain the maintenance of the inverted pattern after the sister *ori* finally segregate and into the cycle of the daughter cells.

### The organisation of the chromosome

Two different organisation patterns of the chromosome of slow-growing *E. coli* have been described. The earlier view is based on FISH analysis of *ori* and *ter* and nucleoid labelling (Bates and Kleckner, 2005; Fisher et al., 2013; Niki et al., 2000). The chromosome is initially organised longitudinally with *ori* and *ter* at opposite extremes of the nucleoid. The *ori* moves towards mid-cell/nucleoid where it is replicated. Sister *ori* then migrate outwards to opposite edges of the nucleoid, which correspond to the quarter positions of the cell. A similar pattern occurs in *Bacillus subtilis* (Wang et al., 2014). The second view, based primarily on snapshot imaging of live cells (Nielsen et al., 2006b, 2006a; Wang et al., 2006, 2005), the chromosome initially adopts a lengthwise (‘transverse’) left-*ori*-right orientation with *ori* at mid-cell and the two chromosomal arms on either side connected by a stretched terminus region. During chromosome replication, the sister *ori* segregate to the cell quarters and the left-*ori*-right pattern is reproduced in each cell half, ensuring inheritance by the daughter cells. Subsequent studies from other groups have reproduced these results (Wiggins et al., 2010; Woldringh et al., 2015) and this transverse (left-*ori*-right or ‘sausage’) model has become the accepted picture of chromosome organisation in *E. coli* during slow growth. However, longitudinal organisation is still relevant as two live-cell studies have found its appearance at faster growth (~1 hour doubling time) with and without overlapping replication (Cass et al., 2016; Youngren et al., 2014).

Indeed, our work shows that a longitudinal organization can occur during slow growth. However, the *ori* are not located at the extreme edge of the nucleoid but close to the outer quarter positions i.e. 25% of the nucleoid mass lies between each *ori* and the closest pole. Thus we refer to it as a mixed or longitudinal-like organsation. This may also explain how the studies referred to above based on imaging loci from each chromosomal arm but not the nucleoid could conclude that the chromosome is organised transversely while others (and this study) support a more longitudinal arrangement.

We also considered the possibility that the different growing conditions of our approach could play a role. In the mother machine cells are continuously fed with fresh media and thereby maintained in steady state growth (Fig. S1D). However, growth on agarose pads should be stable for a few generations so we find this explanation unlikely. A difference in growth media is another potential explanation. However, we found the position distributions remain centered on the quarter position of the nucleoid mass when cells are grown in AB media supplemented with glycerol (Fig. S9) as in the previous studies of the same MG1655 wildtype strain (Nielsen et al., 2006b, 2006a). The same was true using M9 supplemented with glycerol (Fig. S9). However, the distributions are substantially broader than what we found using M9 glucose, which may also contribute to the differing conclusion drawn in the previous studies.

### Quantitative timings of cell cycle events

Our high throughput timelapse approach allows the determination of the entire distribution of steady-state cell cycle event timings and lengths. In particular, we measure a median time of 30 min between *ori* focus duplication and *ter* centralisation and 45 min between *ter* centralisation and stable nucleoid constriction, with these events occurring in this order in 87% of cell cycles (Fig. S3D). These results complement recent studies on the relative timing of replication termination and the onset of cell constriction (Kar et al., 2023; Tiruvadi-Krishnan et al., 2022) and contribute to our understanding of cell cycle progression. The large size of our dataset (tens of thousands of cell cycles) also allows us to quantify the substantial variation between cells. While we have focused on time-based measurements, given that there is a known dependence of chromosome replication initiation on cell size (Levin and Taheri-Araghi, 2019), we also considered the cell length at which cell cycle events occurred (Fig. S1, S3). However, comparing kymograph (based on relative cell age) and demographics (based on cell length) we saw no indication that cell size is a better metric for studying *ter* centralisation. On the other hand, since *ter* centralisation was proposed to be coordinated with its replication (Espéli et al., 2012), we reasoned that a time-based analysis would be most appropriate.

Surprisingly, given the mean 133 min cell cycle duration, we found that 15% of cells were born with two *ori* foci, the majority of which do not initiate replication within their cell cycle (Fig. S1F). This occurs due to the initiation of a second replication (C period) late in the cycle of the mother cell that then continues into the daughter cells. While this observation cautions against the textbook picture of one initiation per cell cycle, studying these ‘outlier’ cell cycles may be informative for the study of cell cycle control (Le Treut et al., 2021; Levin and Taheri-Araghi, 2019).

## Materials and Methods

### Strains, plasmids and media

All strains used in this study are derivatives of E. coli M1655. For imaging origin and terminus, a new strain was constructed by transduction of *glmS::parS_P1_*::kan from strain RM29 (Li et al., 2002; Mercier et al., 2008) and *matP-YPet*::frt::kan::frt from strain RH3 (Männik et al., 2016) into MG1655 (lab collection) respectively. Two plasmids - pFHCP1-CFP and pFHCP1-mTurquoise2, derived from plasmid pFHC2973 (Nielsen et al., 2006b) by deletion of *ygfp-parB_pMT1_*, were used to drive the expression of CFP-ParB_P1_ and mTurquoise2-ParB_P1_ respectively. To create the triple labelled strain IS129, strain RH3 was transduced with *glmS::parS*_P1_::kan after removal of the kanamycin resistance. A detailed list of strains and plasmids used in this study can be found in Table S1 and S2. All experiments, unless otherwise mentioned, were performed at 30°C using M9 minimal media supplemented with 0.2% glucose, 2 mM MgSO_4_, 0.1 mM CaCl_2_. For experiments involving AB minimal media, the recipe and growth conditions of (Nielsen et al., 2006b) were followed. For mother machine experiments, media were supplemented with 0.5 mg/mL BSA (for passivation to reduce cell adhesion to PDMS) and 50 μM IPTG (for induction of mTurquoise2-ParB or CFP-ParB).

### Microscopy

Strains were grown overnight in the respective minimal media without BSA and IPTG. For induction of mTurquoise2-ParB or CFP-ParB for *ori* visualization, 50 μM IPTG was added 3 hours before loading into the mother machine microfluidics device. The microfluidics device was prepared and loaded as described previously (Köhler et al., 2022). The cells in exponential phase were then loaded into the mother machine using a 1 mL syringe. After loading, cells were fed with fresh M9 minimal media supplemented with 0.5 mg/mL BSA and 50 μM IPTG at a rate of 2 μL/min, and data was acquired after 3 hours. Time-lapse images were taken every 5 minutes using a Nikon Eclipse Ti-E with a 100x oil-immersion objective and a HamamatSu Photonics camera. Both phase contrast and fluorescence signals were captured as mentioned above for up to 72 hours. Visualisation of *ori* required blue light excitation (wavelength 436±20 nm), which is known to cause cell cycle arrest (El Najjar et al., 2020). We optimised our imaging settings to avoid this and allow sustained imaging over several days. With these settings, cells divided on average 13% later but we saw no evidence of cell cycle arrest or other defects. The same imaging settings were used for all strains in this study (even for strain IS173, in which mTurquoise2-ParB is not present). The IPTG concentration used for induction of mTurquoise2-ParB or CFP-ParB did not result in a change in growth rate or cell cycle duration under our imaging conditions.

### Image analysis

All analyses were performed using MATLAB. Time-lapse microscopy images were analyzed using our custom-built pipeline called Mothersegger, as previously described (Köhler et al., 2023, 2022). Briefly, time-lapse images acquired using the method described above were saved as TIFF stacks. The pipeline then identifies and isolates individual growth channels, performs background subtraction, and runs segmentation. The segmented data is used to identify cells belonging to the same cell cycle, along with their parent and daughters. A hard cut-off was put in place to discard cell cycles that are less than 10 frames (50 minutes) and greater than 60 frames (300 minutes) for experiments in glucose media. For experiments in glycerol media, the upper limit was raised to 80 frames (400 minutes). After identifying cell cycles, foci detection is performed on relevant fluorescence channels.

### *ori* duplication

To define the *ori* focus duplication, we analyzed complete cell cycles to identify the first frame in which two *ori* foci were detected for the first time. For cells that have an *ori* focus duplication recorded before the first 4 frames, we go back to its mother cell to identify or confirm the correct timing of *ori* focus duplication. In cases where the duplication is recorded in the mother cell, a negative frame number is recorded for *ori* duplication for the daughter cell.

### MatP relocalization

We defined MatP relocalization (centralization) time point as the first of the three consecutive frames (15 min) in which MatP is seen at the mid-cell for the first time in the cell cycle. The mid-cell region is defined as the central 4.8 pixels (0.32 μm) in each cell. This value is identified by analyzing the spread of MatP foci positions in the demograph (Fig. S1H) when it is tightly localized at mid-cell (between 2.48 and 3.01 μm long cells).

### Nucleoid constriction and HU contours

The line profiles of HU-mCherry are obtained by plotting the mean signal along the short axis. A constricted nucleoid is defined by a dip in the middle one third of the HU-mCherry signal greater than the threshold value. The threshold value (0.13) is given by the 95th percentile of the relative depth of the nucleoid signal in new-born cells i.e. the first bin of the plot in Figure S3B. The threshold intensity values defining the contour lines of the HU-mCherry kymographs were chosen such that that 50% or 80% of the HU-mCherry signal was above the threshold, averaged over each cell age bin of the kymograph.

## Supplementary Information

**Figure S1.**
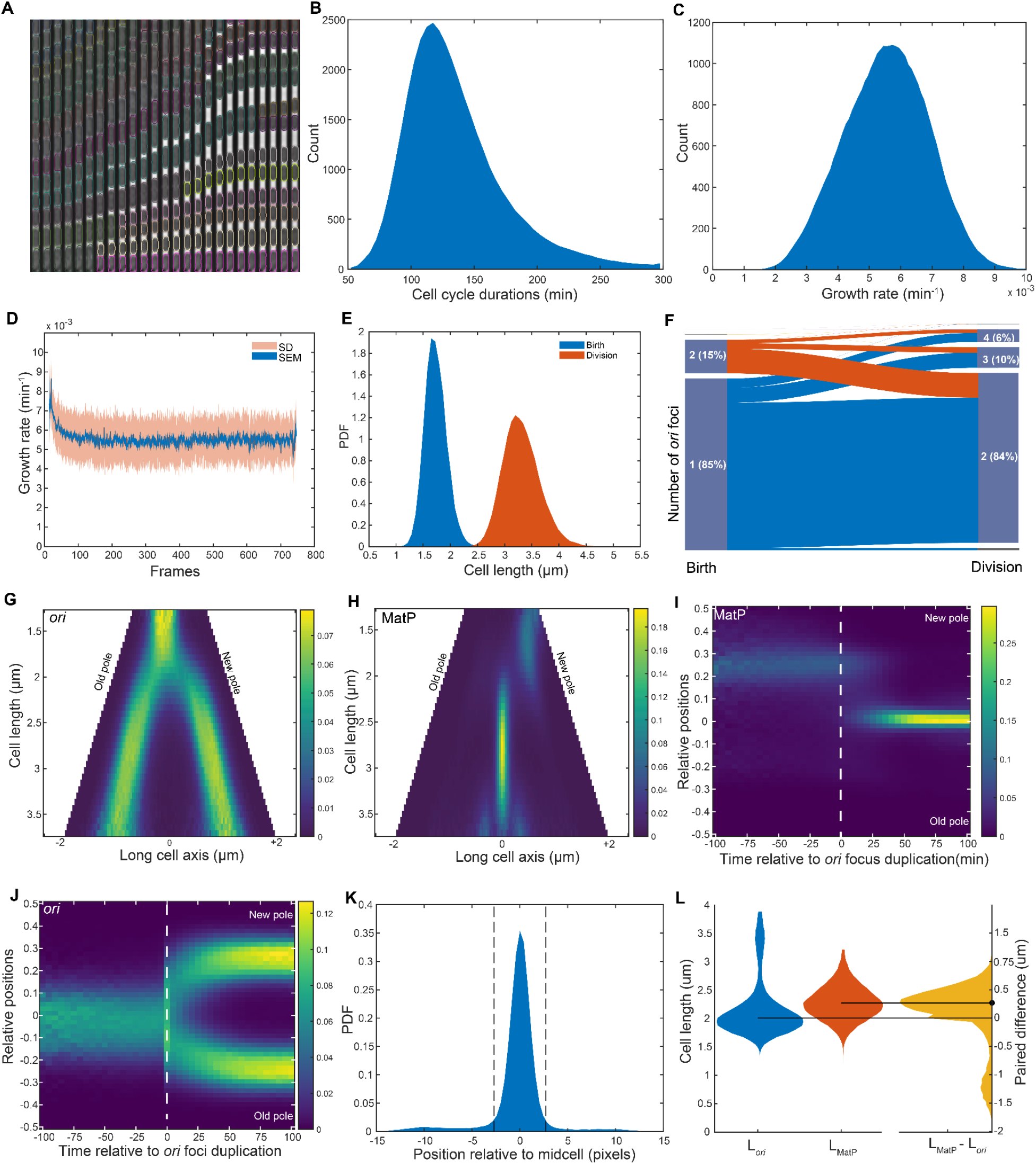
**A.** An example of segmentation and tracking of cells in a growth channel **B.** Histogram of cell cycle durations (mean ± sd = 132.8 ± 39.1 min) for the dataset used in Figure 1 and S1 (n=38066) **C.** Histogram of growth rates for cell cycles ((5.5±1.3)x10^-3^) **D.** Mean cellular growth rate showing stable growth conditions throughout the imaging every 5 minutes **E.** Distribution of birth (1.71 ± 0.2 μm) and division lengths (3.31 ± 0.33 μm) for cells. **F.** Flow diagram showing the number of *ori* foci at birth and division **G.** Demograph of *ori* foci positions along the long axis of cells binned according to cell lengths **H.** Demograph of MatP foci positions along the long axis of cells binned according to cell lengths **I.** Kymograph of MatP foci positions relative to the frame of *ori* foci duplication **J.** Kymograph of *ori* foci positions relative to the frame of *ori* foci duplication **K.** Distribution of MatP foci positions in cells between 2.5 μm and 3 μm. **L.** Distribution of cell lengths at *ori* duplication and MatP centralization along with their paired difference as in Figure 1C. The smaller peak for L_ori_ near 3.5 μm shows that a portion of cells duplicates *ori* twice in a cell cycle (see S1F). The color scale in **I** and **J** is as in Figure 1. The values in the color scale for the demographs **G** and **H** represent the frequency of occurrence of foci positions normalized to the number of cells at each cell length

**Figure S2.**
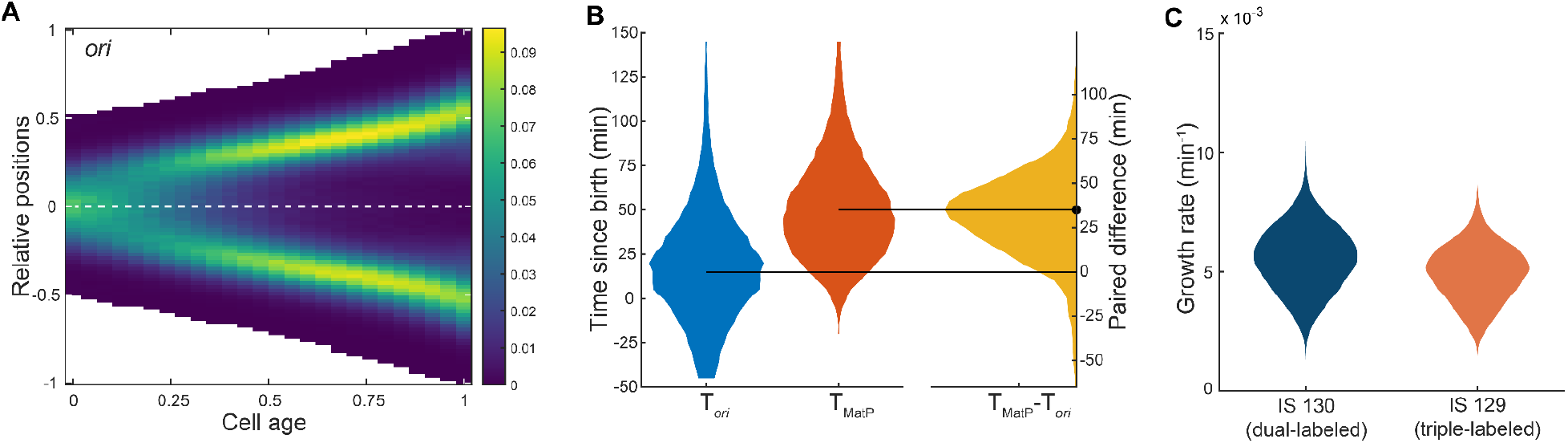
**A.** Average kymograph of *ori* foci positions along the long axis of the triple labeled strain (IS 129) from Figure 2 but with cells oriented randomly, i.e, not according to old-pole and new-pole **B.** Distribution of the time of *ori* focus duplication, T*_ori_* (mean ± sd = 19.1 ± 31.3 min) and MatP, T_MatP_ centralization (50.7 ± 28.0 min) along with the time difference between the two events (31.6 ± 25.2 min) as in Figure 1C **C.** Growth rate distribution of the dual labeled strain IS130 used in Figure 1 ((mean ± sd = (5.5±1.3) x 10^-3^) and triple labeled strain IS129 used in Figure 2 ((4.9±1.2)x10^-3^). The mean doubling times of IS130 and IS 129 are 132.8±39.1 minutes and 142.7±43.5 minutes. The triple labeled strain grows 11% more slowly. Growth rate is calculated using an exponential fit to the cell area for each cell cycle. The color scale in **A** is as in Figure 1.

**Figure S3.**
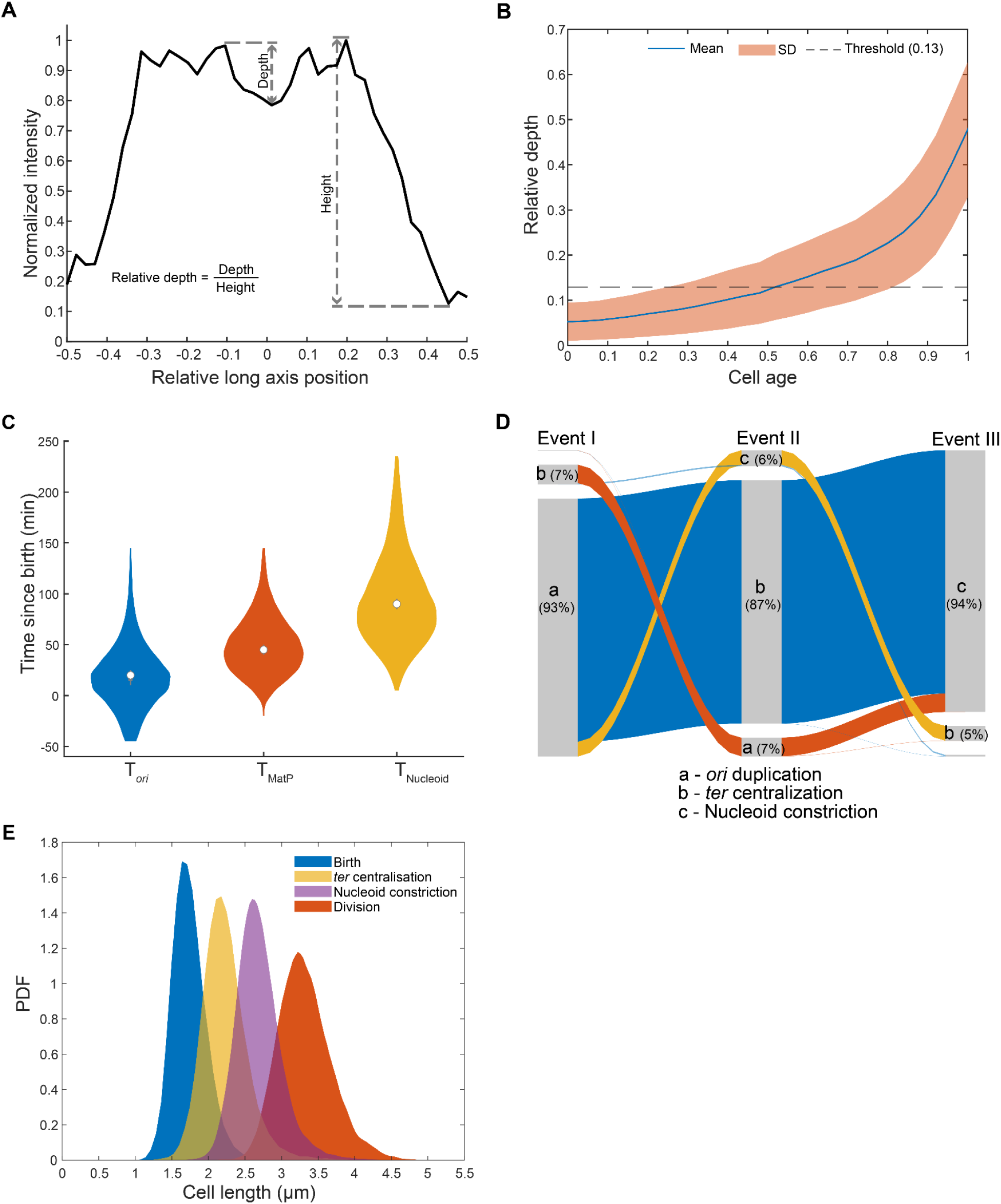
**A**. Line profile from Figure 3A labeled with parameters used for determining nucleoid constriction. **B.** Relative depth of HU-mCherry signal binned according to the cell age (24 bins). The threshold (0.13) for nucleoid constriction is defined as the value at the 95th percentile of the first bin based on the assumptions that no new born cell has a truly constricted nucleoid so that any dips observed are due to random fluctuations. The blue line represents the mean and the shaded region represents standard deviation **C.** Distribution of time of *ori* focus duplication, T*_ori_* (mean ± s.d = 20.0 ± 32.3 min) MatP centralization, T_MatP_ (49.7 ± 28.6 min) and continuous nucleoid constriction, T_Nucleoid_ (95.0 ± 44.3 min). The white dots indicate the mean. The time of stable nucleoid constriction corresponds to the earliest frame starting from the nucleoid has dip greater than the threshold. Paired differences are given in Fig. S2B and Fig. 3B **D.** Order of occurrence of events involving *ori* duplication, *ter* centralization and continuous nucleoid constriction. a, b and c represent the events *ori* duplication, *ter* centralization and nucleoid constriction respectively. The corresponding percentages indicate the proportion of cells in which a particular event was first (Event I), second (Event II) or third (Event III) to occur. **E.** Distribution of cell lengths at which T_MatP_ (2.22 ± 0.30 μm) and T_Nucleoid_ (2.68 ± 0.30 μm) occur in cell cycles along with their birth (1.73 ± 0.22 μm) and division lengths (3.32 ± 0.36 μm).

**Figure S4.**
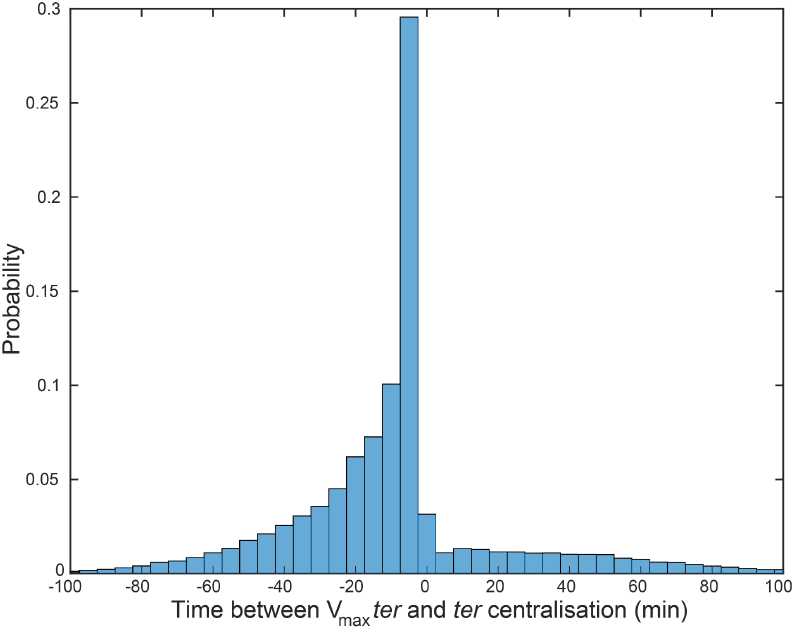
Distribution of time between the frame that has maximum MatP focus velocity (step-wise) and the frame in which MatP centralization occurs. Negative values indicate the maximum MatP velocity occurred before MatP centralisation. Bin width is 5 minutes. Data as in Figure 1, 4.

**Figure S5.**
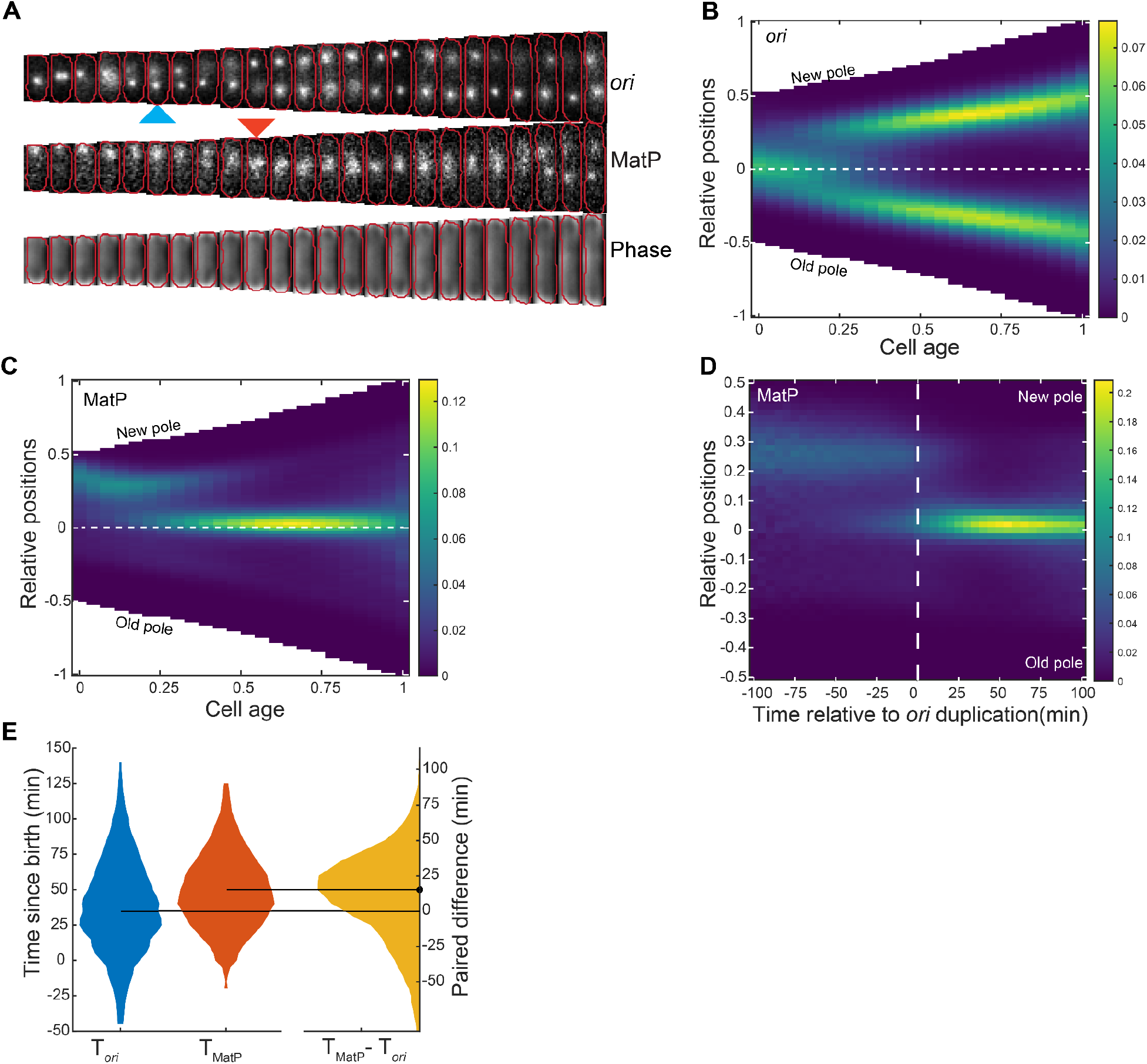
**A.** An example cell cycle of *zapB* strain with *ori* and MatP labeled. The blue arrow indicates *ori* focus duplication and the red arrow indicates MatP relocalization to mid-cell. **B.** Average foci position kymograph of *ori* in a *zapB* strain **C.** Average foci position kymograph of MatP in a *matP*. **D.** Kymograph of MatP foci positions synchronized to *ori* focus duplication. **E.** Distribution of the time of *ori* focus duplication, T*_ori_* (mean ± sd = 38.5 ± 33.7 min) and MatP centralization, T_MatP_ (49.1 ± 26.6 min) along with the time difference between the two events (10.7 ± 31.8 min) similar to Figure 1C. The horizontal lines indicate the median values of 35 and 50 minutes for T*_ori_* and T_MatP_ respectively. The dot indicates the median (15 minutes) of the paired difference T_MatP_ - T*_ori_*. The color scale in **B, C** and **D** is as in Figure 1.

**Figure S6.**
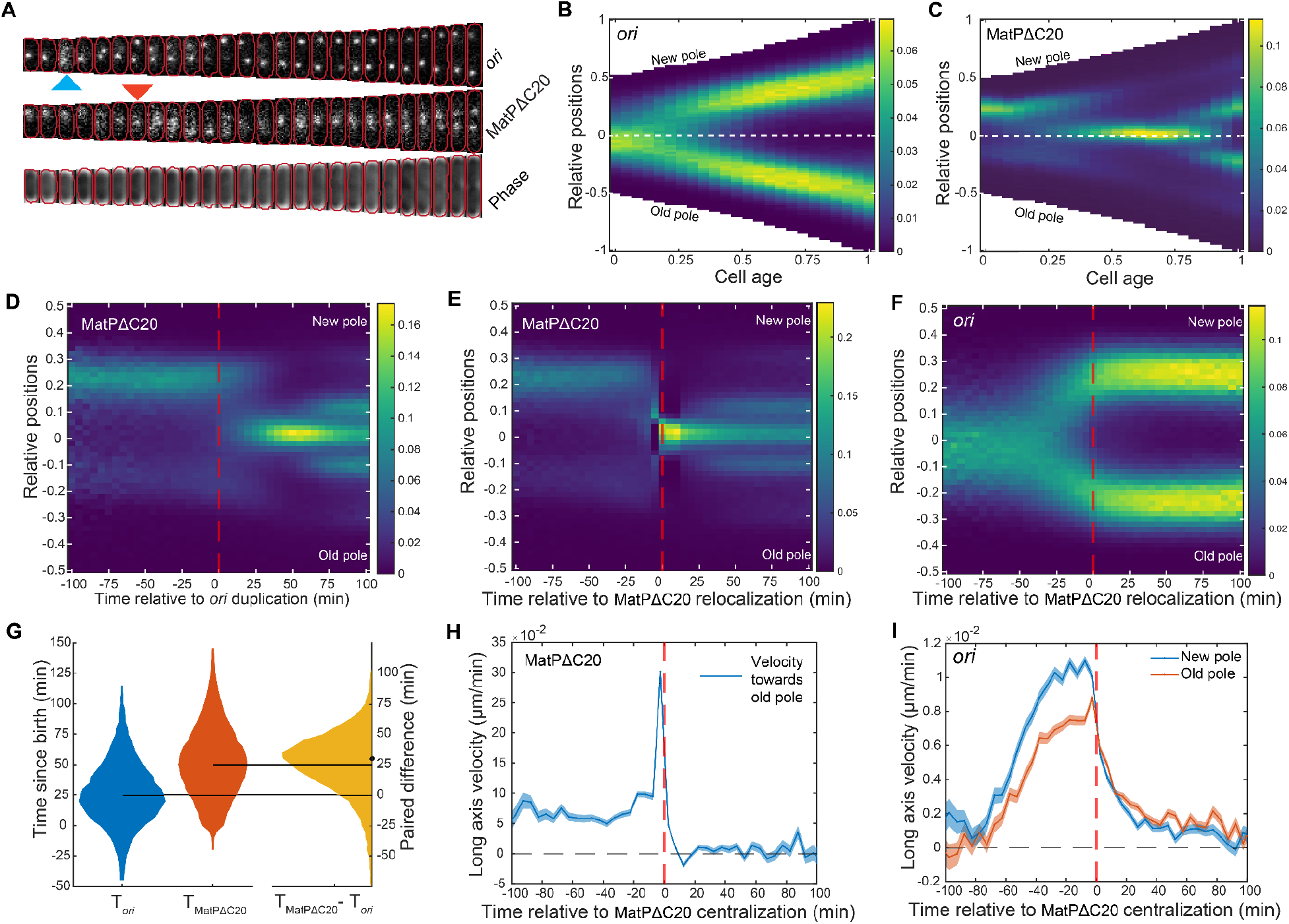
**A.** An example cell cycle of *matPΔC20* strain with *ori* and MatPΔC20 labeled. The blue arrow indicates *ori* focus duplication and the red arrow indicates MatPΔC20 relocalization to mid-cell. **B.** Average foci position kymograph of *ori* in a *matPΔC20* strain **C.** Average foci position kymograph of MatPΔC20 in a *matPΔC20* strain. **D.** Kymograph of MatPΔC20 foci positions synchronized to *ori* focus duplication. **E.** Kymograph of MatPΔC20 foci positions synchronized to MatPΔC20 centralization. **F.** Kymograph of *ori* foci positions relative to MatPΔC20 centralization. **G.** Distribution of the time of *ori* focus duplication, T*_ori_* (mean ± sd = 26.7 ± 26.8 min) and MatPΔC20 centralization, T_MatPΔC20_ (49.1 ± 26.6 min) along with the time difference between the two events (10.7 ± 31.8 min) similar to Figure 1C. The horizontal lines indicate the median values of 35 and 50 minutes for T*_ori_* and T_MatPΔC20_ respectively. The dot indicates the median (15 minutes) of the paired difference T_MatPΔC20_ - T*_ori_*. **G.** Mean velocity of MatPΔC20 focus track towards the old pole relative to MatPΔC20 centralisation. **H.** Mean velocity of *ori* foci tracks towards the nearest pole relative to the time of MatPΔC20 centralisation. Data is from 18202 cell cycles. The color scale in **B, C, D, E** and **F** is as in Figure 1.

**Figure S7.**
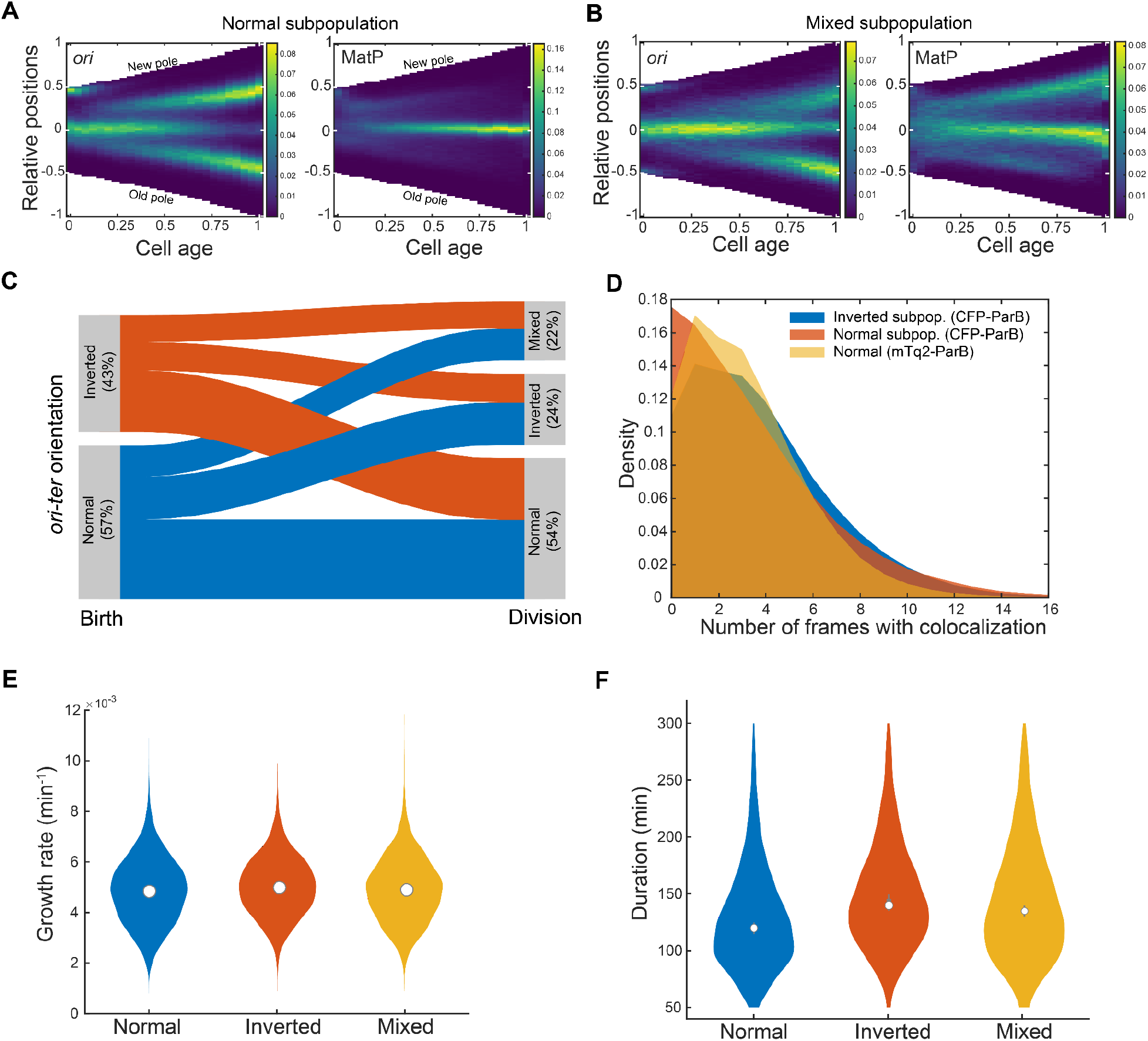
**A.** Foci positions kymographs of cells in the normal subpopulation (n=8269 cells) of CFP-ParB labeled cells. **B.** Foci position kymographs of cells in mixed subpopulation (n=3407 cells). The cells are oriented according to the position of the pole proximal MatP focus on the last frame of the cell cycle. **C.** Flow diagram showing how frequently cells born with normal or inverted *ori-ter* orientation produce daughters with normal, inverted or mixed orientations (n=12384 cells). **D.** Total number of frames in each cell cycle where MatP and *ori* foci are colocalized for the data shown in Figure 6D. The colocalization is defined as frames in which any MatP and *ori* foci appear within 260 pixels. **E.** The distributions of growth rates of the normal (mean ± sd = (4.9±1.3)x10^-3^ min^-1^), inverted ((5.0±1.2)x10^-3^ min^-1^) and mixed ((4.9±1.3)x10^-3^ min^-1^) subpopulations. The white circle indicates the mean. **F.** The distributions of cell cycle durations in normal (129 ± 46 min), inverted (148 ±46 min) and mixed (142 ± 50 min) subpopulations. The white circles indicate the mean. The normal, inverted and mixed subpopulations had 8269, 3774 and 3407 cell cycles respectively. Cell cycles shorter than 50 minutes and longer than 300 minutes were excluded from analysis (see methods). The color scale in **A** and **B** is as in Figure 1.

**Figure S8.**
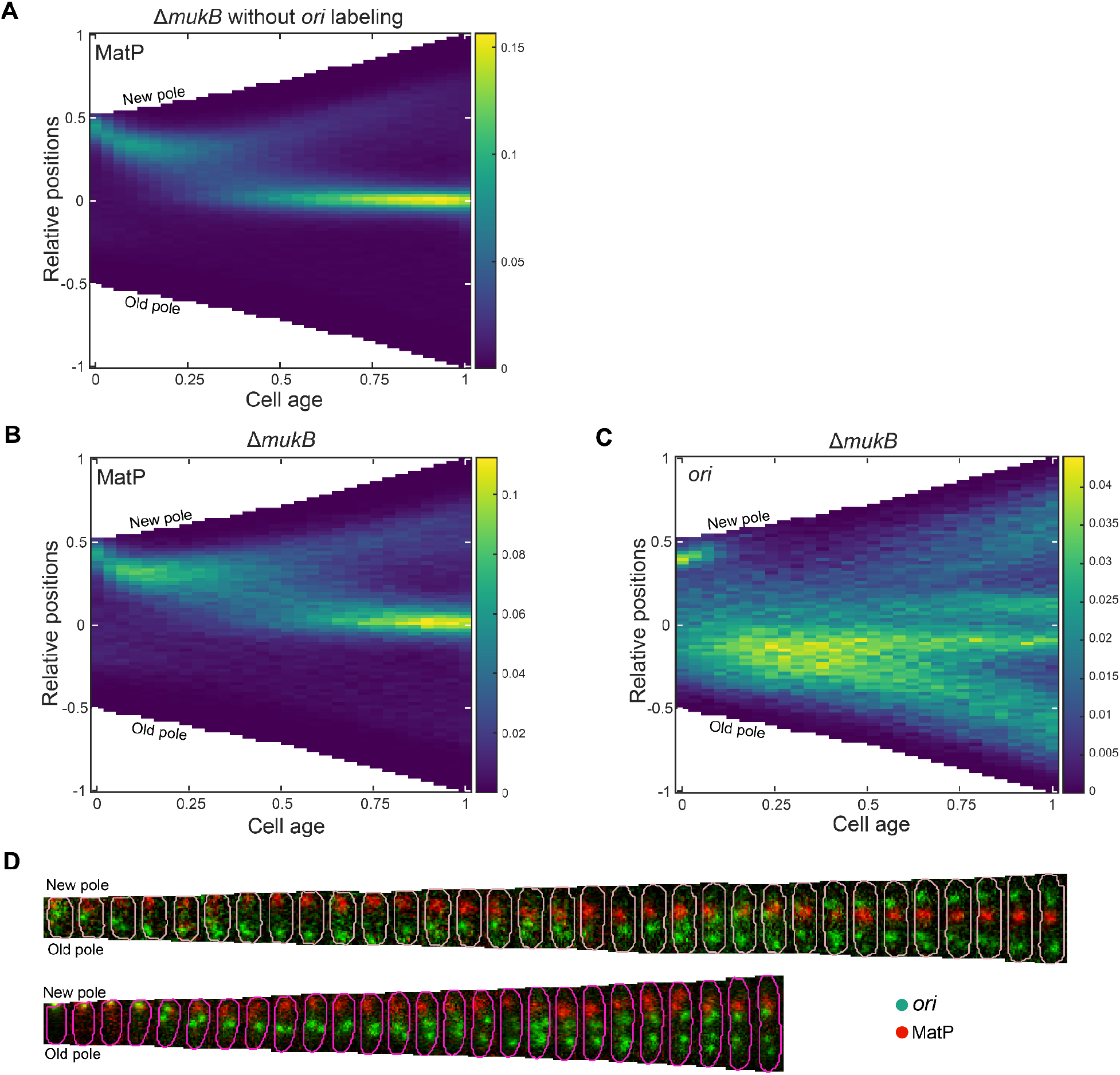
**A.** Foci position kymograph of MatP in a *mukB* strain in a strain without *ori* labelled (n=11062 cell cycles). **B.** and **C.** Foci position kymograph of MatP (**B**) and *ori* (**C**) in a *mukB* strain with *ori* additionally labeled using mTurquoise2-ParB_P1_ (n=3122 cell cycles). Cell cycles producing anucleate daughters (detected by the absence of MatP foci) are not included in the analysis. **D.** Representative cell cycles showing MatP (red) and *ori* (green) dynamics in the *mukB* strain. The color scale in **A, B** and **C** is as in Figure 1.

**Figure S9.**
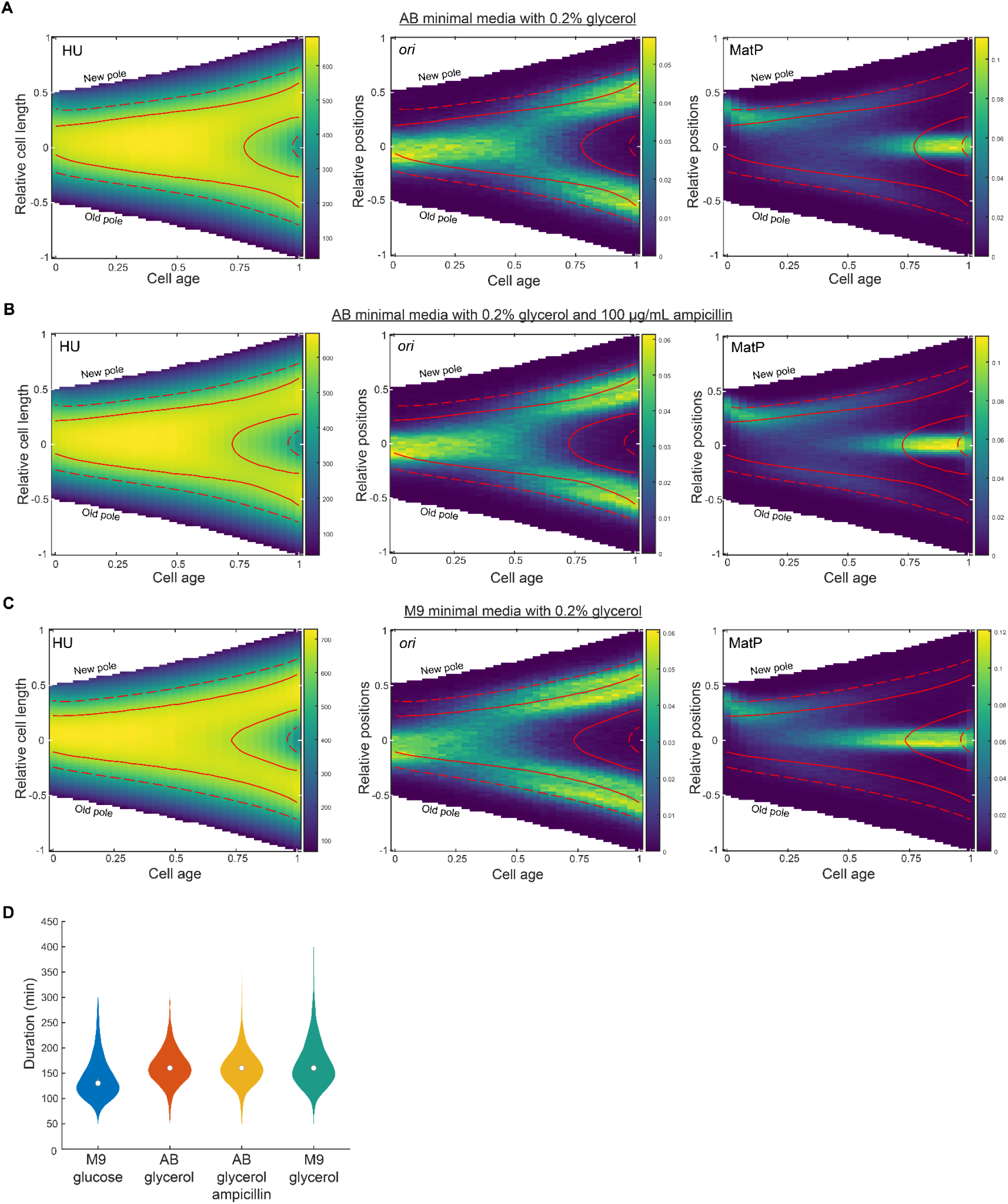
**A.** (Left) Average kymograph of HU-mCherry signal along the long axis of the cell. The solid contour lines represent upper 50 percent and the dashed lines represent upper 80 percent of the total HU-mCherry signal. Average kymograph of *ori* foci positions (middle) and MatP foci positions (right) with contour lines from HU-mCherry grown in AB minimal media with 0.2% glycerol, 1 μg/mL thiamine and 1 μg/mL uracil (n=7281). **B.** Same as **A**, but grown in AB minimal media with 0.2% glycerol, 1 μg/mL thiamine, 1 μg/mL uracil and 100 μg/mL ampicillin (n=8128). **C.** Same as **A, B**, but grown in M9 minimal media supplemented with 0.2% glycerol (n=6973). **D.** Growth rate of IS 129 strain in different media conditions. The white dots indicates mean - M9 minimal media with glucose (same as **Fig. 2**) (mean±s.d = 140.7±41.7 min), AB minimal media (same as **S9A**) (162.8±35.8 min), AB minimal media with ampicillin (same as **S9B**) (161.1±35.9 min) and M9 minimal media with 0.2% glycerol, 2 mM MgSO_4_, 0.1 mM CaCl_2_ (169.6±48.1 min). All experiments except M9 glucose were done at 32°C. The color scales are as in Figure 2.

**Table S1:** Strains used in this study.

| Strain name | Genotype | Source |
| --- | --- | --- |
| IS 111 | MG1655 *matP*-YPet::frt<br>*glmS*::*parS*<sub>P1</sub>::kan + pFHCP1-CFP | This study |
| IS 129 | MG1655 *hupA*-mCherry::frt<br>*matP*-YPet::frt *glmS*::*parS*<sub>P1</sub>::kan +<br>pFHCP1-mTurquoise2 | This study; MG1655<br>*hupA*-mCherry::frt<br>*matP*-YPet::frt::kan::frt received<br>from Jaan Männik (Männik et al.,<br>2016) |
| IS 130 | MG1655 *matP*-YPet::frt<br>*glmS*::*parS*<sub>P1</sub>::kan +<br>pFHCP1-mTurquoise2 | This study |
| IS 146 | MG1655 *matP*Δ*C20*-YPet::frt<br>*glmS*::*parS*<sub>P1</sub>::kan +<br>pFHCP1-mTurquoise2 | This study |
| IS 161 | MG1655 Δ*zapB*::frt::cat::frt<br>*matP*-YPet::frt *glmS*::*parS*<sub>P1</sub>::kan +<br>pFHCP1-mTurquoise2 | This study |
| IS 173 | MG1655 Δ*mukB*::frt::kan::frt<br>*matP*-YPet::frt<br>*glmS*::*parS*<sub>P1</sub>::frt::cat::frt | This study |
| IS 174 | MG1655 Δ*mukB*::frt::kan::frt<br>*matP*-YPet::frt<br>*glmS*::*parS*<sub>P1</sub>::frt::cat::frt +<br>pFHCP1-mTurquoise2 | This study |

**Table S2:** Plasmids used in this study.

| Plasmid name | Use | Source |
| --- | --- | --- |
| pFHCP1-CFP | Used to express CFP-ParB<sub>P1</sub><br>needed to visualize *ori*. Inducible<br>with IPTG. | This study; Derived from plasmid<br>pFHC2973 (Nielsen et al., 2006b) |
| pFHCP1-mTurquoise2 | Used to express<br>mTurquoise2-ParB<sub>P1</sub> needed to<br>visualize *ori*. Inducible with IPTG. | This study; Made by replacement of<br>CFP with mTurquoise2 in<br>pFHCP1-CFP |

**Table S3:**
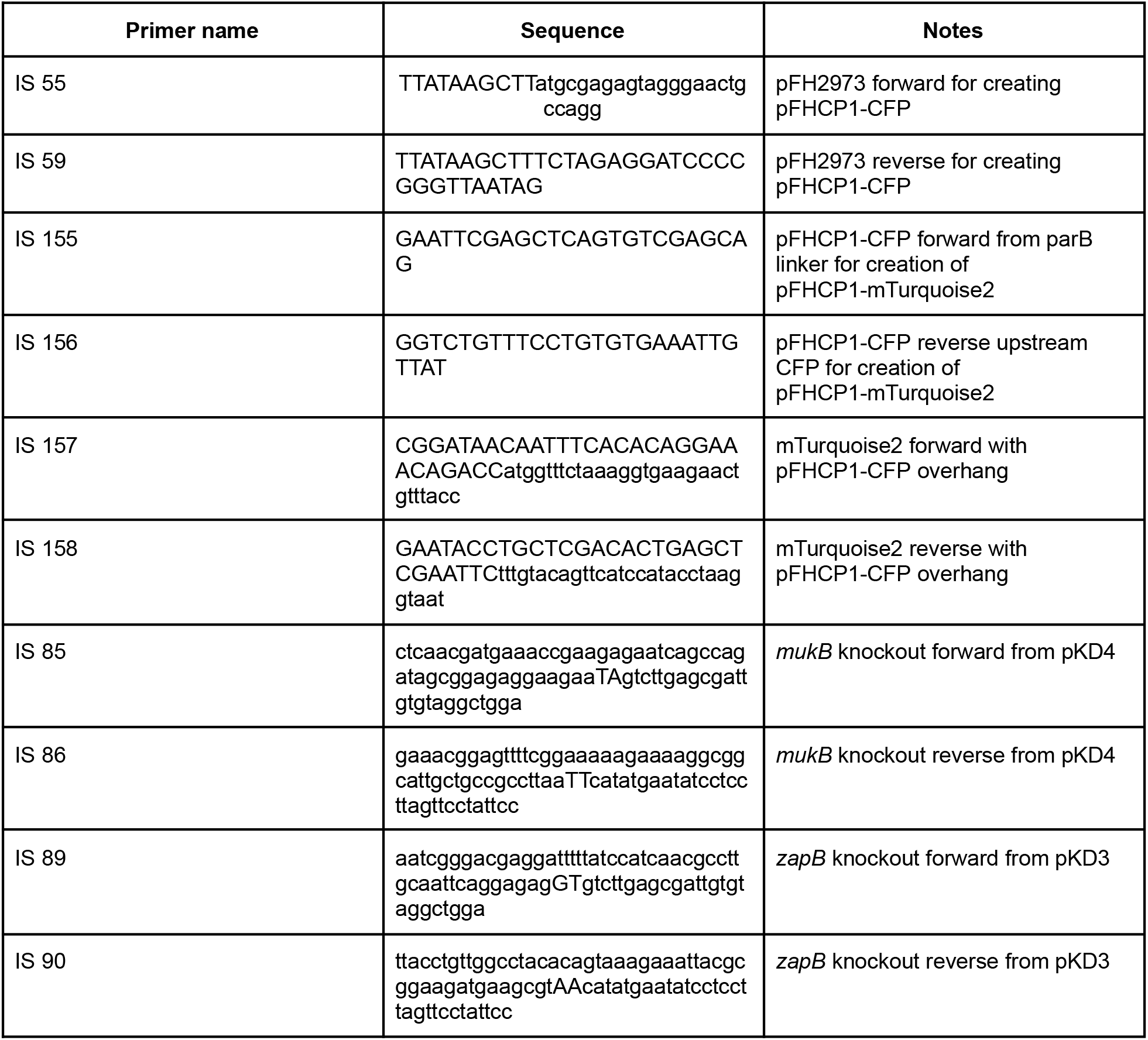
Primers used in this study.

## References

Badrinarayanan A, Lesterlin C, Reyes-Lamothe R, Sherratt DJ. 2012. The escherichia coli SMC complex, MukBEF, shapes nucleoid organization independently of DNA replication. J Bacteriol 194:4669–4676. doi: 10.1128/JB.00957-12

Bailey MW, Bisicchia P, Warren BT, Sherratt DJ, Männik J. 2014. Evidence for divisome localization mechanisms independent of the Min system and SlmA in Escherichia coli. PLoS Genet 10:e1004504. doi: 10.1371/journal.pgen.1004504

Bates D, Kleckner N. 2005. Chromosome and Replisome Dynamics in E. coli: Loss of Sister Cohesion Triggers Global Chromosome Movement and Mediates Chromosome Segregation. Cell 121:899–911. doi: 10.1016/j.cell.2005.04.013

Böhm K, Meyer F, Rhomberg A, Kalinowski J, Donovan C, Bramkamp M. 2017. Novel Chromosome Organization Pattern in Actinomycetales—Overlapping Replication Cycles Combined with Diploidy. mBio 8:e00511–17. doi: 10.1128/mBio.00511-17

Cass JA, Kuwada NJ, Traxler B, Wiggins PA. 2016. Escherichia coli Chromosomal Loci Segregate from Midcell with Universal Dynamics. Biophys J 110:2597–2609. doi: 10.1016/j.bpj.2016.04.046

Crozat E, Tardin C, Salhi M, Rousseau P, Lablaine A, Bertoni T, Holcman D, Sclavi B, Cicuta P, Cornet F. 2020. Post-replicative pairing of sister ter regions in Escherichia coli involves multiple activities of MatP. Nat Commun 11:1–12. doi: 10.1038/s41467-020-17606-6

Danilova O, Reyes-Lamothe R, Pinskaya M, Sherratt DJ, Possoz C. 2007. MukB colocalizes with the oriC region and is required for organization of the two Escherichia coli chromosome arms into separate cell halves. Mol Microbiol 65:1485–92. doi: 10.1111/j.1365-2958.2007.05881.x

David A, Demarre G, Muresan L, Paly E, Barre F-X, Possoz C. 2014. The Two Cis-Acting Sites, parS1 and oriC1, Contribute to the 10ngitudinal Organisation of Vibrio cholerae Chromosome I. PloS Genet 10:e1004448. doi: 10.1371/journal.pgen.1004448

Donachie WD. 1968. Relationship between Cell Size and Time of Initiation of DNA Replication. Nature 219:1077–1079. doi: 10.1038/2191077a0

Dupaigne P, Tonthat NK, Espéli O, Whitfill T, Boccard F, Schumacher MA. 2012. Molecular basis for a protein-mediated DNA-bridging mechanism that functions in condensation of the E. coli chromosome. Mol Cell 48:560–71. doi: 10.1016/j.molcel.2012.09.009

El Najjar N, van Teeseling MCF, Mayer B, Hermann S, Thanbichler M, Graumann PL. 2020. Bacterial cell growth is arrested by violet and blue, but not yellow light excitation during fluorescence microscopy. BMC Mol Cell Biol 21:35. doi: 10.1186/s12860-020-00277-y

Espéli O, Borne R, Dupaigne P, Thiel A, Gigant E, Mercier R, Boccard F. 2012. A MatP-divisome interaction coordinates chromosome segregation with cell division in E. coli. EMBO J 31:3198–211. doi: 10.1038/emboj.2012.128

Fekete RA, Chattoraj DK. 2004. A cis-acting sequence involved in chromosome segregation in Escherichia coli. Mol Microbiol 55:175–183. doi: 10.1111/j.1365-2958.2004.04392.x

Fisher JK, Bourniquel A, Witz G, Weiner B, Prentiss M, Kleckner N. 2013. Four-dimensional imaging of E. coli nucleoid organization and dynamics in living cells. Cell 153:882–895. doi: 10.1016/j.cell.2013.04.006

Hadizadeh Yazdi N, Guet CC, Johnson RC, Marko JF. 2012. Variation of the folding and dynamics of the Escherichia coli chromosome with growth conditions. Mol Microbiol 86:1318–1333. doi: 10.1111/mmi.12071

Hofmann A, Mäkelä J, Sherratt DJ, Heermann D, Murray SM. 2019. Self-organised segregation of bacterial chromosomal origins. eLife 8:e46564. doi: 10.7554/eLife.46564

Jun S, Mulder B. 2006. Entropy-driven spatial organization of highly confined polymers: Lessons for the bacterial chromosome. Proc Natl Acad Sci 103:12388–12393. doi: 10.1073/pnas.0605305103

Kar P, Tiruvadi-Krishnan S, Männik J, Männik J, Amir A. 2023. Using conditional independence tests to elucidate causal links in cell cycle regulation in Escherichia coli. Proc Natl Acad Sci 12¤:e2214796120. doi: 10.1073/pnas.2214796120

Köhler R, Kaganovitch E, Murray SM. 2022. High-throughput imaging and quantitative analysis uncovers the nature of plasmid positioning by ParABS. eLife 11. doi: 10.7554/eLife.78743

Köhler R, Sadhir I, Murray SM. 2023. ✶Track: Inferred counting and tracking of replicating DNA loci. Biophys J in press.

Lau IF, Filipe SR, Søballe B, Økstad O-A, Barre F-X, Sherratt DJ. 2003. Spatial and temporal organization of replicating Escherichia coli chromosomes. Mol Microbiol 49:731–743. doi: 10.1046/j.1365-2958.2003.03640.x

Le Treut G, Si F, Li D, Jun S. 2021. Quantitative Examination of Five Stochastic Cell-Cycle and Cell-Size Control Models for Escherichia coli and Bacillus subtilis. Front Microbiol 12.

Lemonnier M, Bouet J-Y, Libante V, Lane D. 2000. Disruption of the F plasmid partition complex in vivo by partition protein SopA. Mol Microbiol 38:493–503. doi: 10.1046/j.1365-2958.2000.02101.x

Levin PA, Taheri-Araghi S. 2019. One is Nothing without the Other: Theoretical and Empirical Analysis of Cell Growth and Cell Cycle Progression. J Mol Biol 431:2061–2067. doi: 10.1016/j.jmb.2019.04.004

Li Y, Sergueev K, Austin S. 2002. The segregation of the Escherichia coli origin and terminus of replication. Mol Microbiol 46:985–995. doi: 10.1046/j.1365-2958.2002.03234.x

Lioy VS, Cournac A, Marbouty M, Duigou S, Mozziconacci J, Espéli O, Boccard F, Koszul R. 2018. Multiscale Structuring of the E. coli Chromosome by Nucleoid-Associated and Condensin Proteins. Cell 172:771–783.e18. doi: 10.1016/j.cell.2017.12.027

Mäkelä J, Uphoff S, Sherratt DJ. 2021. Nonrandom segregation of sister chromosomes by *Escherichia coli* MukBEF. Proc Natl Acad Sci 118:e2022078118. doi: 10.1073/pnas.2022078118

Männik J, Bailey MW. 2015. Spatial coordination between chromosomes and cell division proteins in Escherichia coli. Front Microbiol 6:306. doi: 10.3389/fmicb.2015.00306

Männik J, Castillo DE, Yang D, Siopsis G, Männik J. 2016. The role of MatP, ZapA and ZapB in chromosomal organization and dynamics in *Escherichia coli*. Nucleic Acids Res 44:gkv1484. doi: 10.1093/nar/gkv1484

Mercier R, Petit M-A, Schbath S, Robin S, El Karoui M, Boccard F, Espéli O. 2008. The MatP/matS site-specific system organizes the terminus region of the E. coli chromosome into a macrodomain. Cell 135:475–85. doi: 10.1016/j.cell.2008.08.031

Murray SM, Sourjik V. 2017. Self-organization and positioning of bacterial protein clusters. Nat Phys 13:1006–1013. doi: 10.1038/nphys4155

Nicolas E, Upton AL, Uphoff S, Henry O, Badrinarayanan A, Sherratt DJ. 2014. The SMC complex MukBEF recruits topoisomerase IV to the origin of replication region in live Escherichia coli. mBio 5:e01001–13. doi: 10.1128/mBio.01001-13

Nielsen HJ, Li Y, Youngren B, Hansen FG, Austin S. 2006a. Progressive segregation of the Escherichia coli chromosome. Mol Microbiol 61:383–393. doi: 10.1111/j.1365-2958.2006.05245.x

Nielsen HJ, Ottesen JR, Youngren B, Austin SJ, Hansen FG. 2006b. The *Escherichia coli* chromosome is organized with the left and right chromosome arms in separate cell halves. Mol Microbiol 62:331–338. doi: 10.1111/j.1365-2958.2006.05346.x

Niki H, Hiraga S. 1998. Polar localization of the replication origin and terminus in Escherichia coli nucleoids during chromosome partitioning. Genes Dev 12:1036–1045.

Niki H, Yamaichi Y, Hiraga S. 2000. Dynamic organization of chromosomal DNA in *Escherichia coli*. Genes Dev 14:212–23. doi: 10.1101/GAD.14.2.212

Nolivos S, Upton AL, Badrinarayanan A, Müller J, Zawadzka K, Wiktor J, Gill A, Arciszewska LK, Nicolas E, Sherratt DJ. 2016. MatP regulates the coordinated action of topoisomerase IV and MukBEF in chromosome segregation. Nat Commun 7:10466. doi: 10.1038/ncomms10466

Stouf M, Meile J-C, Cornet F. 2013. FtsK actively segregates sister chromosomes in *Escherichia coli*. Proc Natl Acad Sci 110:11157–11162. doi: 10.1073/pnas.1304080110

Teleman AA, Graumann PL, Lin DC-H, Grossman AD, Losick R. 1998. Chromosome arrangement within a bacterium. Curr Biol 8:1102–1109. doi: 10.1016/S0960-9822(98)70464-6

Tiruvadi-Krishnan S, Männik J, Kar P, Lin J, Amir A, Männik J. 2022. Coupling between DNA replication, segregation, and the onset of constriction in Escherichia coli. Cell Rep 38:110539. doi: 10.1016/j.celrep.2022.110539

Vallet-Gely I, Boccard F. 2013. Chromosomal Organization and Segregation in Pseudomonas aeruginosa. PLOS Genet 9:e1003492. doi: 10.1371/journal.pgen.1003492

Viollier PH, Thanbichler M, McGrath PT, West L, Meewan M, McAdams HH, Shapiro L. 2004. Rapid and sequential movement of individual chromosomal loci to specific subcellular locations during bacterial DNA replication. Proc Natl Acad Sci U S A 101:9257–62. doi: 10.1073/pnas.0402606101

Wang P, Robert L, Pelletier J, Dang WL, Taddei F, Wright A, Jun S. 2010. Robust growth of Escherichia coli. Curr Biol 20:1099–103. doi: 10.1016/j.cub.2010.04.045

Wang X, Liu X, Possoz C, Sherratt DJ. 2006. The two Escherichia coli chromosome arms locate to separate cell halves. Genes Dev 20:1727–1731. doi: 10.1101/gad.388406

Wang X, Montero Lopis P, Rudner DZ. 2014. Bacillus subtilis chromosome organization oscillates between two distinct patterns. Proc Natl Acad Sci U S A 111:12877–12882. doi: 10.1073/pnas.1407461111

Wang X, Possoz C, Sherratt DJ. 2005. Dancing around the divisome: asymmetric chromosome segregation in Escherichia coli. Genes Dev 19:2367–77. doi: 10.1101/gad.345305

Wiggins PA, Cheveralls KC, Martin JS, Lintner R, Kondev J. 2010. Strong intranucleoid interactions organize the Escherichia coli chromosome into a nucleoid filament. Proc Natl Acad Sci U S A 107:4991–5. doi: 10.1073/pnas.0912062107

Woldringh CL, Hansen FG, Vischer NOE, Atlung T. 2015. Segregation of chromosome arms in growing and non-growing Escherichia coli cells. Front Microbiol 6.

Youngren B, Nielsen HJ, Jun S, Austin S. 2014. The multifork Escherichia coli chromosome is a self-duplicating and self-segregating thermodynamic ring polymer. Genes Dev 28:71–84. doi: 10.1101/gad.231050.113

